# Extended family with an inherited pathogenic variant in polymerase delta provides strong evidence for recessive effect of proofreading deficiency in human cells

**DOI:** 10.1101/2022.07.20.500591

**Authors:** Maria A. Andrianova, Vladimir B. Seplyarskiy, Mariona Terradas, Ana Beatriz Sánchez-Heras, Pilar Mur, José Luis Soto, Gemma Aiza, Fyodor A. Kondrashov, Alexey S. Kondrashov, Georgii A. Bazykin, Laura Valle

## Abstract

Mutational processes in germline and in somatic cells are vastly different, and it remains unclear how the same genetic background affects somatic and transmissible mutations. Here, we estimate the impact of an inherited pathogenic variant in the exonuclease domain of polymerase delta (Polδ) on somatic and germline mutational processes and cancer development. In germline cells and in non-cancer somatic cells, the *POLD1* L474P variant increases the mutation burden only slightly, contributing ∼11.8% and ∼14.7% of mutations respectively, although it strongly distorts the mutational spectra. By contrast, tumors developed by carriers of inherited pathogenic variants in *POLD1* harbor a DNA rearrangement that results in a homozygous state of the pathogenic variant, leading to an extremely high mutation rate. Thus, mutations in both alleles of *POLD1* gene are required for strong increase in mutation rate suggesting recessiveness of Poldδ proofreading. These results show a similar role of Polδ in germline and somatic replication, and, together with previous findings, illustrate the important differences between Polδ and Polε in the disruption of their replication fidelity.

## INTRODUCTION

Changes in the DNA sequence can arise due to errors during replication or due to DNA damage caused by intracellular or environmental factors. Mutation rate and patterns are drastically different between somatic and germline cells (Heredia-Genestar et al. 2020; Moore et al. 2020; Milholland et al. 2017), likely due to different exposures to external and internal mutagens or differences in proficiency of DNA repair. The relative contribution of replicative and non-replicative errors remains a matter of debate for both somatic and germline mutations, although this interpretation has been questioned recently (de Manuel et al. 2022). In cancer cells, most mutations result from exposure to exogenous and endogenous factors (Moore et al. 2020; Abascal et al. 2021). A dominant contribution of replication errors is only observed in cancer cells deficient in the proofreading of replicative polymerases or in replication-associated mismatch repair (Meier et al. 2018; Degasperi et al. 2022; Alexandrov et al. 2013; Haradhvala et al. 2016, 2018).

Cells with deficiencies in systems controlling replication fidelity have a high potential for the study of the impact of replicative mutations: first, they have unique mutational spectra (Haradhvala et al. 2018), and second, the accumulated mutations depend on the number of cell divisions. The strong conservation of replication mechanisms in eukaryotes (Fukui 2010; Kunkel 2009) suggests that replication deficiencies likely affect both germline and soma in a similar way. In particular, inherited mutations that affect the proofreading function of the main replicative polymerases, i.e. polymerases epsilon and delta, increase the mutation rate of normal human cells including spermatocytes, although the effect varies between polymerases and among different pathogenic variants (Robinson et al. 2021). To understand the impact of replication errors and the effect of a particular inherited variant that causes decreased replication fidelity on somatic and germline mutations simultaneously, and to distinguish the effect between male and female germline, we studied the mutation rate and spectra in an extended family harboring *POLD1* c.1421T>C p.(Leu474Pro) (from herein on, L474P). The effect of this variant on the proofreading activity of Pol**δ** and its association with cancer predisposition is supported by multiple pieces of evidence and may be currently classified as likely pathogenic (Supplemental Note, Supplemental Fig. S1). To estimate the genetic effect of this mutation in human somatic cells and minimize the possible influence of the microenvironment, the tissue structure and other factors, we cultured fibroblast single-cell colonies and measured the increase of mutation rate within a controlled experimental system. Altogether, we showed that L474P contributes a similar proportion of mutations to germline cells and fibroblasts, only slightly increasing the overall mutation burden but significantly disturbing mutational spectra.

Inherited pathogenic mutations affecting the proofreading activity of Polδ cause a multi-organ tumor predisposition syndrome, where colorectal adenomatous polyposis, colorectal and endometrial cancers are the most prevalent phenotypic features (Palles et al. 2022). Cancer development in the carriers is thought to be associated with an increased somatic mutation rate. However, as it was shown previously (Robinson et al. 2021) and confirmed in our study, heterozygous inactivation of Polδ proofreading leads to just a modest increase in mutation rate. Studies in yeasts and mice suggested a recessive effect of Polδ proofreading inactivation both in terms of mutation rate (Simon et al. 1991; Morrison et al. 1993; Zhou et al. 2021; Goldsby et al. 2002) and phenotype (Goldsby et al. 2002, 2001; Albertson et al. 2009). A deeper investigation of the cancer samples developed by patients harboring inherited *POLD1* cancer-predisposing variants uncovered that, unlike polymerase epsilon (Polε), deficiency of proofreading in both copies of *POLD1* gene is likely required for *POLD1*-driven human carcinogenesis, making applicable the classic Knudson’s two-hit hypothesis for tumor suppressor and DNA repair genes.

## RESULTS

### Extended family with the inherited *POLD1* L474P pathogenic variant

In the family investigated in this study, *POLD1* L474P was identified in eight members, three of whom had been diagnosed with colorectal cancer (ages at cancer diagnosis: 23-50), and one with endometrial cancer at age 58. Most L474P carriers were diagnosed with gastrointestinal polyps, some of them with a clear attenuated adenomatous polyposis phenotype (10-100 adenomas) (Fig. 1A, Supplemental Table S1). It is intuitive to assume that the inherited *POLD1* L474P genotype increases the risk of cancer due to a higher mutation rate in somatic tissues caused by a polymerase proofreading defect. A mutator phenotype was observed when the homologous residue was substituted in the haploid strain of *Saccharomyces cerevisiae* (Murphy et al. 2006) or *Schizosaccharomyces pombe* (Supplemental Fig. S1). However, the mutagenic effect of this variant in heterozygotic state in human normal tissues has been recently shown to be modest (Robinson et al. 2021).

**Figure 1.**
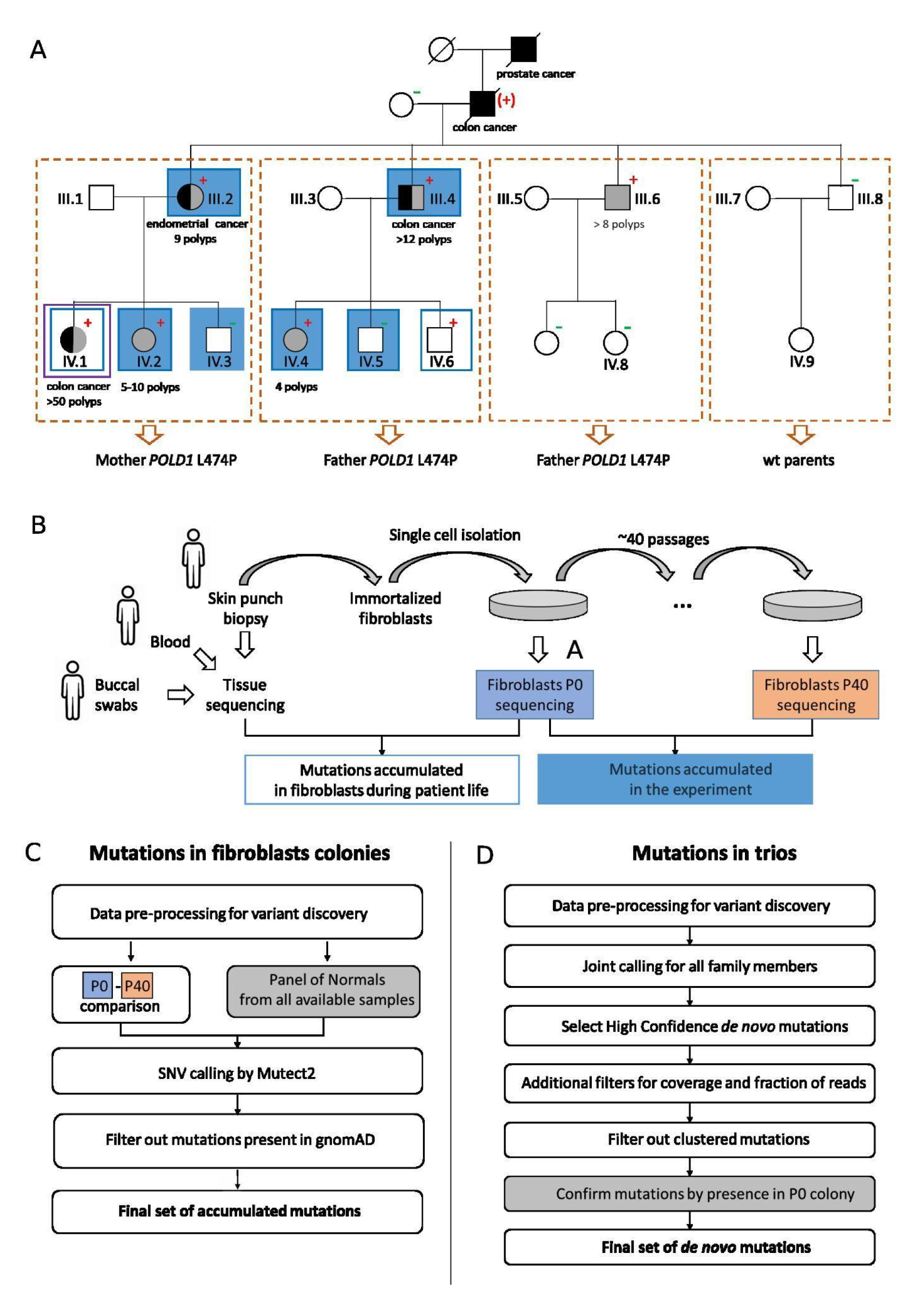
Family, sample description, and study pipeline. **A,** Family pedigree. Individuals diagnosed with cancer are marked in black, and with polyps, in gray. The tumor types and number of polyps are indicated below the corresponding individual symbol. Detailed tumor phenotypes and ages at diagnosis are shown in Supplemental Table S1. Blue rectangles mark the individuals from whom skin biopsies for single-cell fibroblast colonies were obtained, filled rectangles represent colonies that were used for the mutation accumulation experiments. The violet rectangle marks the individual with a sequenced tumor sample. Sequenced sub-families are marked in orange with individual IDs shown for the members that were sequenced. Plus and minus signs mark the carrier status for *POLD1* L474P, and a plus sign in parentheses (+) indicates an obligate carrier. **B,** Scheme of the experiment performed on immortalized fibroblasts from family members for the assessment of somatic mutation accumulation. **C,** Algorithm used for calling somatic mutations accumulated during the growth of the cell lines. **D,** Algorithm used for calling germline *de novo* mutations.

### Influence of inherited *POLD1* L474P on somatic mutagenesis

To study the effect of *POLD1 L474P* on somatic mutation rate, we designed a controlled experiment for six fibroblast colonies (for individuals III.2, IV.2, IV.3, III.4, IV.4 and IV.5, Fig. 1A-filled blue rectangles). The single-cell derived colony for each individual was sequenced at the initial time point. After this, the cell lines were grown for ∼40 passages after single cell isolation (Supplemental Table S2), and genome sequencing was performed on the DNA obtained from the cultured cells at the endpoints of the experiment (Fig. 1B, Methods). The detection of mutations accumulated during the passages was performed using a standard procedure for somatic single nucleotide calling (Fig. 1C, Methods). While the number of mutations accumulated during the experiment in the fibroblasts harboring *POLD1* L474P was higher than that in the wild-type fibroblasts, for most samples this increase was relatively minor (Fig. 2A). For three out of four carriers (III.2, III.4, IV.2), the effect for indels was slightly stronger than for SNVs, however, the opposite was observed for IV.4 (Fig. 2B). Insertions and deletions were dominated by single T insertions in homopolymer tracts, similarly to previous results (Robinson et al. 2021), and 2-bp deletions in long repeats (Supplemental Fig. S2). Although the effect on the overall mutation rate was minor, the *POLD1* L474P carrier status was associated with a radical shift in the mutational spectrum. *De novo* extraction and decomposition of mutational signatures in the endpoint fibroblasts from the four carriers and the two non-carriers of the *POLD1* L474P heterozygous pathogenic variant yielded six COSMIC signatures. Refitting the mutational spectrums to this fixed number of signatures in each sample revealed a highly significant presence of the Polδ proofreading deficiency signature SBS10c in all four *POLD1* L474P carriers but not in non-carriers (Fig. 2C, Supplemental Table S3). In fact, the excess of mutations in *POLD1* L474P fibroblasts may be mostly attributed to SBS10c (Fig. 2D).

**Figure 2.**
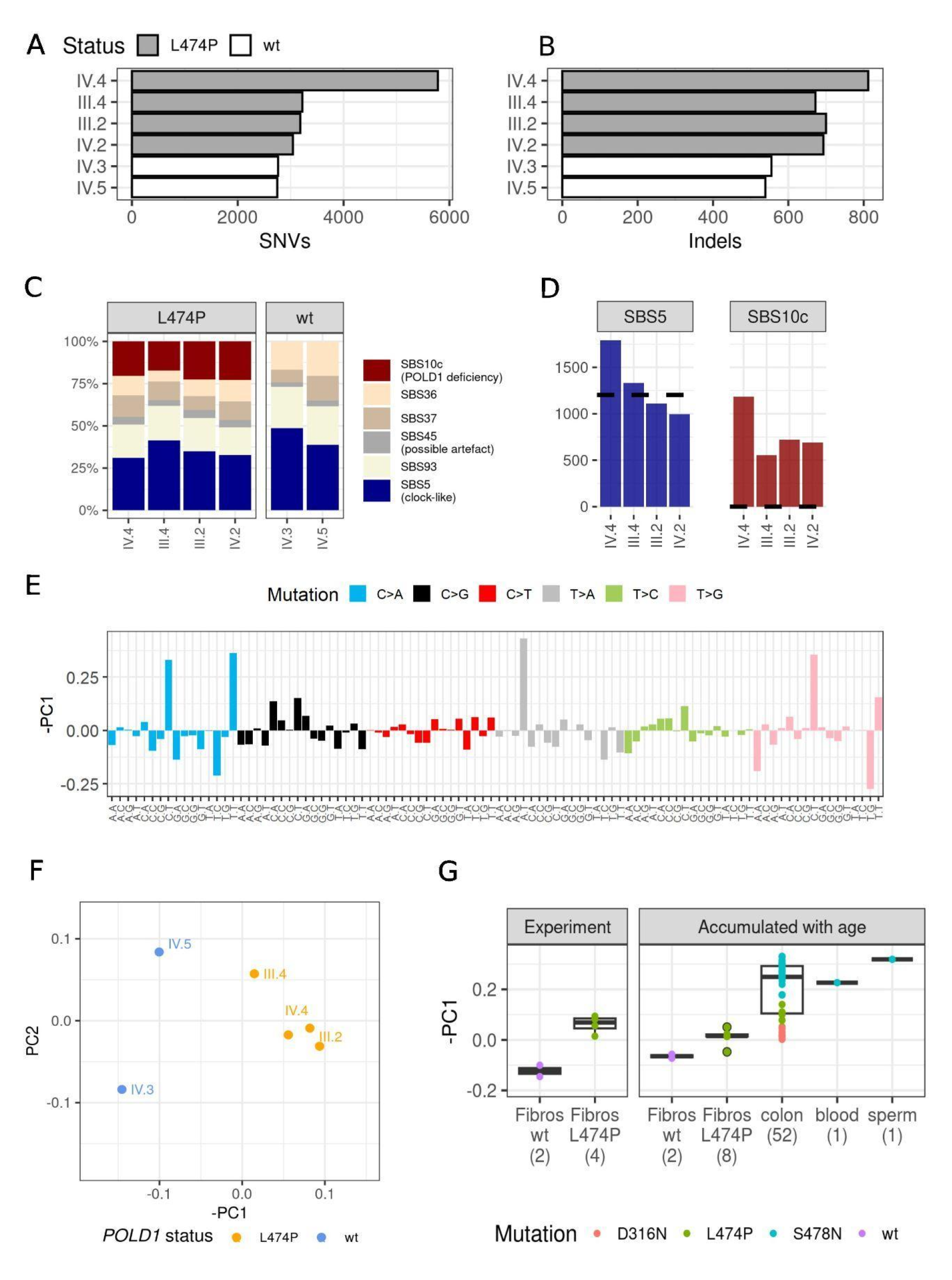
Somatic mutation burden and spectrum. **A-B,** Number of SNVs (**A**) and indels (**B**) accumulated during the experiment in the cultured fibroblasts of carriers and non-carriers of *POLD1* L474P. **C,** Proportion of mutations attributed to the different COSMIC signatures accumulated in the cultured fibroblasts. *De novo* signatures were extracted and decomposed to COSMIC signatures using SigProfilerExtractor with subsequent refitting using SigFit. **D,** Excess of mutations corresponding to the different mutational signatures in the cultured fibroblasts of *POLD1* L474P heterozygous carriers. The black dashed lines mark the mean number of mutations attributable to the corresponding signature in wild-type fibroblasts. **E,** Loadings of mutational contexts in –PC1. **F,** PCA analysis of 96-context mutational spectra of mutations accumulated in the cultured fibroblasts. **G,** Values of the PC1 component in wt and L474P fibroblasts colonies from the mutation accumulation experiment and in other normal tissues obtained from heterozygous carriers of *POLD1* pathogenic variants: skin fibroblasts (sequenced in this study) and other tissues (Robinson et al. 2021). The different colors correspond to different *POLD1* pathogenic variants. Numbers in the parenthesis indicate the number of analyzed samples.

To orthogonally assess the effect of *POLD1* L474P on mutational spectrum, we calculated the 96-context mutational spectra for the fibroblasts with and without *POLD1* L474P, and applied principal component analysis (PCA) to these spectra (Fig. 2F). PCA clearly separated L474P carriers from non-carriers: PC1 explained ∼48% of the variance in the mutational spectrum across samples, and reflected the experimentally obtained difference in spectra between *POLD1* mutated and non-mutated individuals (Fig. 2E). Comparison of the principal components with COSMIC mutational signatures showed that PC1 had the strongest correlation with SBS10c (cosine similarity = 0.52). PC1 also separated previously published samples of somatic tissues from individuals with inherited pathogenic variants in *POLD1* (Fig. 2G). Moreover, the values of PC1 differed between samples with different pathogenic variants in *POLD1:* S478N showed the highest PC1 values, and D316N, the lowest. This is consistent with previous findings that had shown that *POLD1* S478N had the strongest effect on mutational burden, while this effect was much lower for D316N (Robinson et al. 2021).

Similarly to the previous study (Robinson et al. 2021) we also searched for the footprint of Polδ proofreading deficiency in mutations accumulated in tissues over the lifetime of the individual. For this, we analyzed mutational patterns in single-cell derived colonies obtained from immortalized fibroblasts (origin: skin punches) from eight family members (six carriers of *POLD1* L474P and two non-carriers) (blue rectangles in Fig. 1A). Among the mutations accumulated in the skin fibroblasts of the individuals over their lifetime, an estimated 9705 on average were caused by ultraviolet (UV) radiation. Such mutations are manifested by the COSMIC single base substitution signature 7 (SBS7) and a high proportion of CC>TT double substitutions (Supplemental Fig. S3). Interestingly, the number of identified UV-induced mutations was higher than previously reported for skin fibroblasts (Saini et al. 2021) and more in line with the numbers reported for melanocytes (Tang et al. 2020). The number of observed mutations correlates with the individual age at the time of biopsy (Supplemental Fig. S4).

We observed the SBS10c signature, corresponding to Polδ proofreading deficiency, in all carriers of L474P. Unexpectedly, mutations attributed to SBS10c, although in a much smaller number, were also identified in non-carriers. A plausible explanation to this observation could be misattribution due to the refitting procedure to a fixed subset of signatures (see Methods). In fact, simulations confirm that imperfection of signature decomposition could identify the presence of non-existent signatures (Supplemental Figure S5). An orthogonal approach using PCA also confirms the presence of Polδ proofreading deficiency signature (increased values of -PC1) in carriers of L474P (Fig. 2G). The separation from non-carriers was however more fuzzy compared to experimental data, in line with the results of signature analysis and suggesting a noisier structure of the data.

### Influence of inherited *POLD1* L474P on germline mutagenesis

Next, we studied the impact of *POLD1* L474P on mutation patterns in the germline. For this, we sequenced four sub-families from the studied family, each including two parents and one to three children, comprising a total of eight offspring (Fig. 1A, Supplemental Table S1). In two sub-families, *POLD1* L474P was carried by the father; in one sub-family, by the mother; and in the remaining sub-family, both parents had wild-type *POLD1*. *De novo* mutations in each offspring were called using the standard GATK4 pipeline with additional filters (Methods), focused on identifying mutations present in the offspring but absent in both parents (Fig. 1D). The number of called *de novo* mutations per individual varied between 24 and 88 (Fig. 3A).

**Figure 3.**
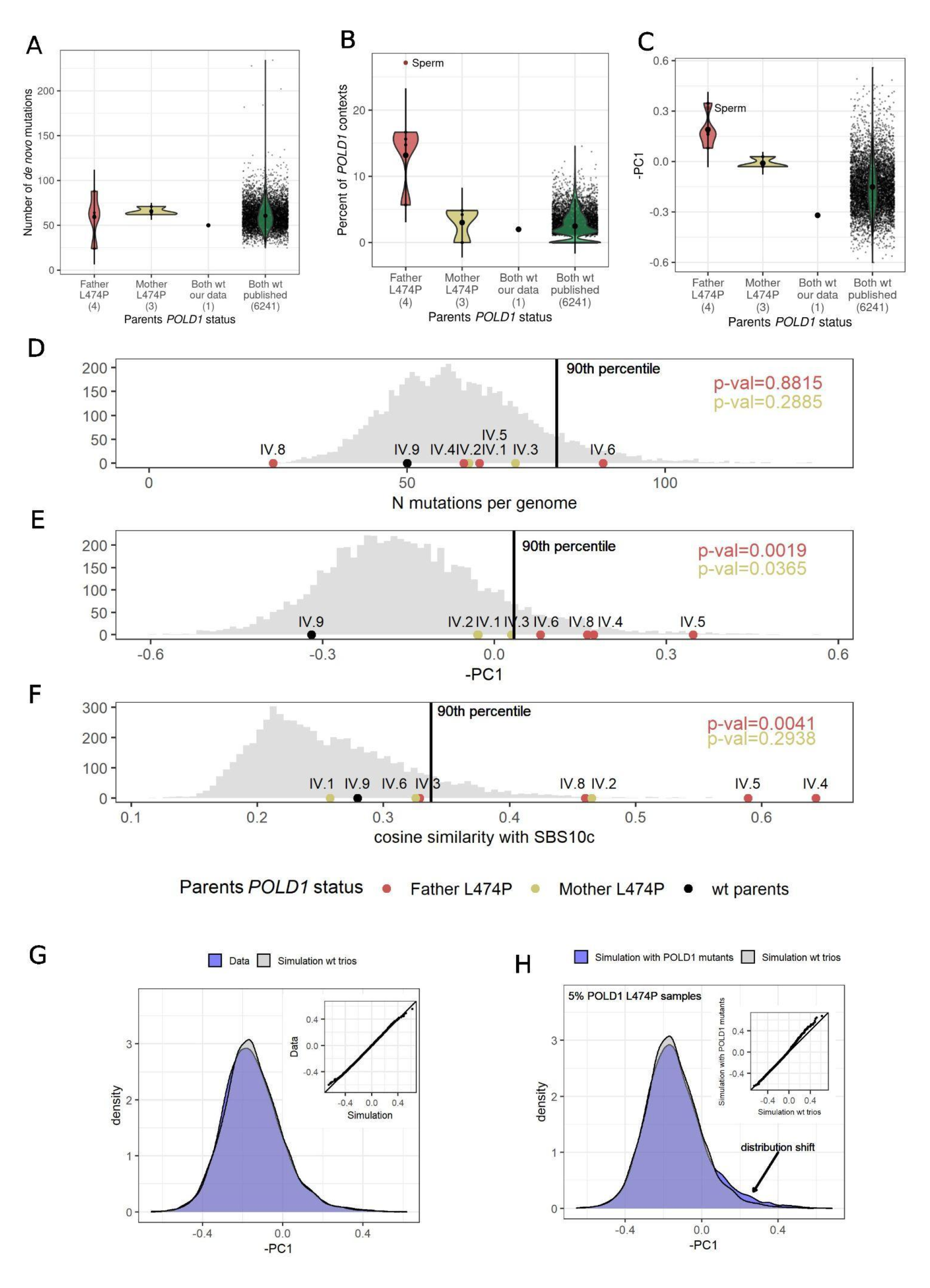
Germline mutation burden and spectrum. **A,** Number of *de novo* mutations in offspring of parents with wild-type or mutated *POLD1.* **B,** Proportion of mutated *POLD1*-specific tri-nucleotide contexts in *de novo* mutations in offspring of parents with wild-type or mutated *POLD1.* **C,** Value of -PC1 in *de novo* mutations in offspring of parents with wild-type or mutated *POLD1.* In **B** and **C**, the red dot (sperm) corresponds to the sperm sample from an individual with a inherited *POLD1* S478N variant previously published (Robinson et al. 2021). In **A-C** numbers in parentheses indicate the number of analyzed samples. **D-F,** Representation of the number of mutations (**D**), -PC1 values (**E**), and cosine similarity with SBS10c signature (**F**) in sequenced trios from the family compared to the distribution of the corresponding parameter in publicly available trios (Halldorsson et al. 2019; An et al. 2018). The p-value for Kolmogorov-Smirnov test of public trios against fathers-carriers of *POLD1* variant is shown in red, and against mothers-carriers of *POLD1* variant is shown in khaki. **G,** Distribution of the -PC1 values in published and simulated *de novo* mutation spectra. **H,** Distribution of -PC1 values in simulations of *de novo* mutations with a mixture of offspring of *POLD1* L474P fathers.

Theoretically, *POLD1* L474P should affect the number of replication errors. However, based on the obtained fibroblast data, we expected this effect to be minor, with a ∼20% increase in the overall number of mutations in the offspring due to the *POLD1* L474P genotype. Because of the ∼10-fold higher number of cell divisions in spermatocytes compared to oocytes (Gao et al. 2019), one would also expect a larger increase in the mutation rate in male gametes. Our limited data showed no significant effect of *POLD1* L474P on the burden of *de novo* mutations, when either the father or the mother carried the variant (Fig. 3A). We next performed an analysis of mutational spectra, which would be better powered, because *POLD1* L474P produces mutations in very specific contexts and its spectrum is well described by four main mutation types: Cp<u>C</u>pT>A, Tp<u>C</u>pT>A, Ap<u>T</u>pT>A and Cp<u>T</u>pT>G (Fig. 2E). The proportion of *de novo* mutations in these contexts among all 96 contexts estimated from previously published trios (Halldorsson et al. 2019; An et al. 2018) was ∼2.4%. Given this and the total number of observed *de novo* mutations per offspring of fathers with *POLD1* L474P variant, we expected a total of 5.7 mutations in these contexts for four offspring. Instead, we observed 28 such mutations (11.8% of all mutations), suggesting a ∼4.9 fold increase (rate ratio test p-value = 6.619e-05, Fig. 3B). These observations indicate that in spermatocytes, the fraction of mutations caused by replicative errors introduced by *POLD1* L474P is comparable to that observed in fibroblasts (14.7% in soma and 11.8% in germline). By contrast, in the offspring of *POLD1* L474P mothers, the fraction of mutations in the four contexts mostly affected by *POLD1* L474P was practically the same as in the wild-type trios (Fig. 3B, Supplemental Fig. S6).

Consistently, the projection of mutational spectrum of *de novo* mutations in the offspring of *POLD1* L474P fathers to -PC1 showed a significantly higher value compared to the offspring of mothers with *POLD1* L474P, and to the offspring of wild-type parents (KS test p-value = 0.0019) (Fig. 3C and 3E). Altogether, our analyses demonstrate that the presence of a mutator allele could be more easily detectable from changes in the mutational spectra than from an increase in the overall mutation burden (Fig. 3D-F). In particular, the offspring of fathers harboring *POLD1* L474P could not be distinguished from the offspring of wild-type parents based on the number of *de novo* mutations (Fig. 3D). However, *de novo* mutations in *POLD1*- specific mutational contexts were clearly enriched in the offspring of *POLD1* L474P fathers compared to the offspring of wild-type parents (Fig. 3E-F). If an additive model of mutational processes is assumed, this observation might seem counterintuitive. However, even in that scenario, a small number of mutations originating from an additional mutagenic process may result in a shift in the mutational spectrum if the spectrum of the additional mutational process strongly differs from the background processes, even though the absolute number of those mutations can be low (simulations supporting this hypothesis are shown in Supplemental Fig. S7).

Recently, pathogenic variants in known DNA repair genes were shown to contribute to germline hypermutation in sequenced human trios (Kaplanis et al. 2022). We wondered if the offspring of parents with deficient Polδ proofreading activity contribute to the public datasets of sequenced families. To address this question, we mapped onto our PCA coordinates the mutational spectra of 6,233 offspring belonging to 4,638 families with assumedly wild-type *POLD1* status. Although the mean value of -PC1 in this sample was significantly lower than that in the offspring of *POLD1* L474P fathers, we found that 138 (2.2%) of the offspring had –PC1 above this value (Fig. 3C). The observed proportion of samples with a high –PC1 value could indicate the presence of undetected Polδ deficiency in parents of these trios; alternatively, it could reflect stochasticity in low count data. To distinguish between these alternatives, we generated an artificial dataset of trios by randomly sampling mutations according to their fractions in the spectra of *de novo* mutations in offspring of wild-type parents. Among these simulated trios, 2.0% of the offspring had values of -PC1 above the mean value observed in the five offspring of the two *POLD1* L474P fathers studied here. The presence of such cases in our simulated dataset indicates that a high value of -PC1 in some of the trios could be obtained by chance, and a comparable proportion of such cases in the data and in the simulation suggests that either *POLD1* mutated fathers are absent in the dataset or their fraction is smaller than the resolution of this method. To estimate the fraction of fathers with *POLD1* L474P that would detectably change the distribution of -PC1 values in a dataset, we generated an additional synthetic cohort by mixing simulated offspring with mutations sampled from the *de novo* spectrum of wild-type parents with small fractions of simulated offspring with mutations sampled from the spectra of offspring of *POLD1* L474P fathers. For the 1% admixture, the proportion of samples with -PC1 above the mean observed in the offspring of *POLD1* L474P fathers was 2.5%, similar to the simulation with wild-type parents only. For the 5% admixture, the distribution had a pronounced shift to higher -PC1 values, and the corresponding proportion was 3.98% (a 1.8-fold increase compared to the wild-type simulation) (Fig. 3H). We can thus conclude that the presence of a mutation rate modifier is detectable in a sample of trios based on the heterogeneity in the mutational spectrum in the offspring if the fraction of parents carrying the modifying mutation is substantial (e.g. ∼5% for *POLD1* L474P).

Sequencing more than one child in a family should provide more power to detect families with deficiency in Polδ proofreading, as it is unlikely to obtain high values of -PC1 in multiple offspring by chance alone. By simulating multiple offspring per family with wild-type parents, we estimated that only 0.16% of the quartets with two sequenced siblings, and 0.04% of the quintets with three sequenced siblings, are expected to have the -PC1 value averaged between siblings higher than the mean -PC1 value in the offspring of fathers harboring *POLD1* L474P (Supplemental Fig. S8). This confirms that sequencing more than one child per family decreases the rate of false positive prediction of a mutation rate modifier presence in the father, thus increasing the power of this approach. Among the 4,638 analyzed families, 1,555 were quartets with two sequenced siblings. Among them, the observed fraction of families with increased -PC1 value averaged between siblings (0.19%) was similar to the 0.16% estimated from the simulation, again arguing against a detectable contribution of families with Polδ deficiency to the studied dataset.

As mentioned above, our results indicate that the presence of an additional mutagenic process has more effect on the mutational spectrum of *de novo* mutations than the overall number of mutations (Supplemental Fig. S7), and deviation from the spectrum of wild-type trios can potentially be used to detect a presence of mutators in the population. This observation can be generalized to any germline mutagenic process. Application of our method to different repair pathways characterized by unique spectra in synthetic datasets suggested that only processes affecting at least 3-7% of a population could be thus detected (Supplemental Fig. S9). Such high frequencies of mutators for rare and highly severe varions such as Polδ or Polε proofreading deficiency in natural populations seem unrealistic. Potential application of this method to more common mutagenic processes needs further investigation.

### Biallelic deficincy of Polδ proofreading is required for hypermutability

Inherited heterozygous mutations in the exonuclease domain of *POLE* or *POLD1* cause PPAP; a highly penetrant cancer predisposition syndrome characterized by numerous polyps in the colon and an increased risk of colorectal cancer, endometrial cancer, and other tumor types (Church 2014). *POLD1* L474P is classified as pathogenic following the ACMG/AMP variant classification guidelines for PPAP (Mur et al. 2020) (Supplemental note), and, as expected, co-segregates with colorectal cancer, endometrial cancer and gastrointestinal benign tumors in the studied family (Fig. 1A). The proofreading function of *POLE* and *POLD1* was previously proposed to be haploinsufficient (The CORGI Consortium et al. 2013; Bellido et al. 2016). However, unlike *POLE* pathogenic variants, which strongly affect germline and somatic mutation rates even in heterozygous state (Robinson et al. 2021), we and others (Robinson et al. 2021) found only a minor effect of heterozygous *POLD1* L474P on the mutation rate in non-tumoral tissues. To better understand how such a weak mutation rate modifier can drive a highly penetrant cancer phenotype (Valle et al. 2014; Bellido et al. 2016; Ferrer-Avargues et al. 2017; Palles et al. 2022), we sequenced a tumor sample from the family with *POLD1* L474P (Fig. 1A; colorectal cancer developed by individual IV.1). In contrast to cultured fibroblasts, the sequenced tumor was characterized by an ultrahypermutable phenotype, with 226 mutations per megabase (Fig. 4A). The mutational spectrum of the tumor also differed from the spectrum identified in the cultured fibroblasts of *POLD1* L474P carriers, and in phenotypically normal crypts from carriers of different inherited *POLD1* pathogenic variants sequenced in another study (Robinson et al. 2021). Specifically, it was enriched in C>A mutations (∼54% of observed mutations), particularly in Tp<u>C</u>pA and Tp<u>C</u>pT contexts (Fig. 4B). This mutational spectrum perfectly matched SBS10d (cosine similarity = 0.96) found in hypermutable polyps from carriers of inherited *POLD1* S478N (Robinson et al. 2021). We also sequenced and analyzed a colorectal cancer developed by a carrier of the pathogenic variant *POLD1* D316H, which affects a catalytic site of the Polδ exonuclease. This tumor was also characterized by a high mutation rate (96.4 mutations/Mb) and prevalence of SBS10d (Fig. 4C). Thus, two tumors have extremely high mutation rate and share the shift in mutational spectrum with a previously published hypermutable polyp from an individual with inherited *POLD1* S478N (Robinson et al. 2021), suggesting a mutational process different from the mild mutagenesis observed in normal tissues of individuals with heterozygous *POLD1* mutations. The two analyzed tumors were microsatellite stable and had normal expression of mismatch repair (MMR) proteins MLH1, MSH2, MSH6 and PMS2, excluding inactivation of this DNA repair system. In addition, tumors with dysfunctional MMR and an error-prone version of Polδ usually have spectra that correspond to mutational signature SBS20 (Alexandrov et al. 2013; Hao et al. 2016), which was not observed here.

**Figure 4.**
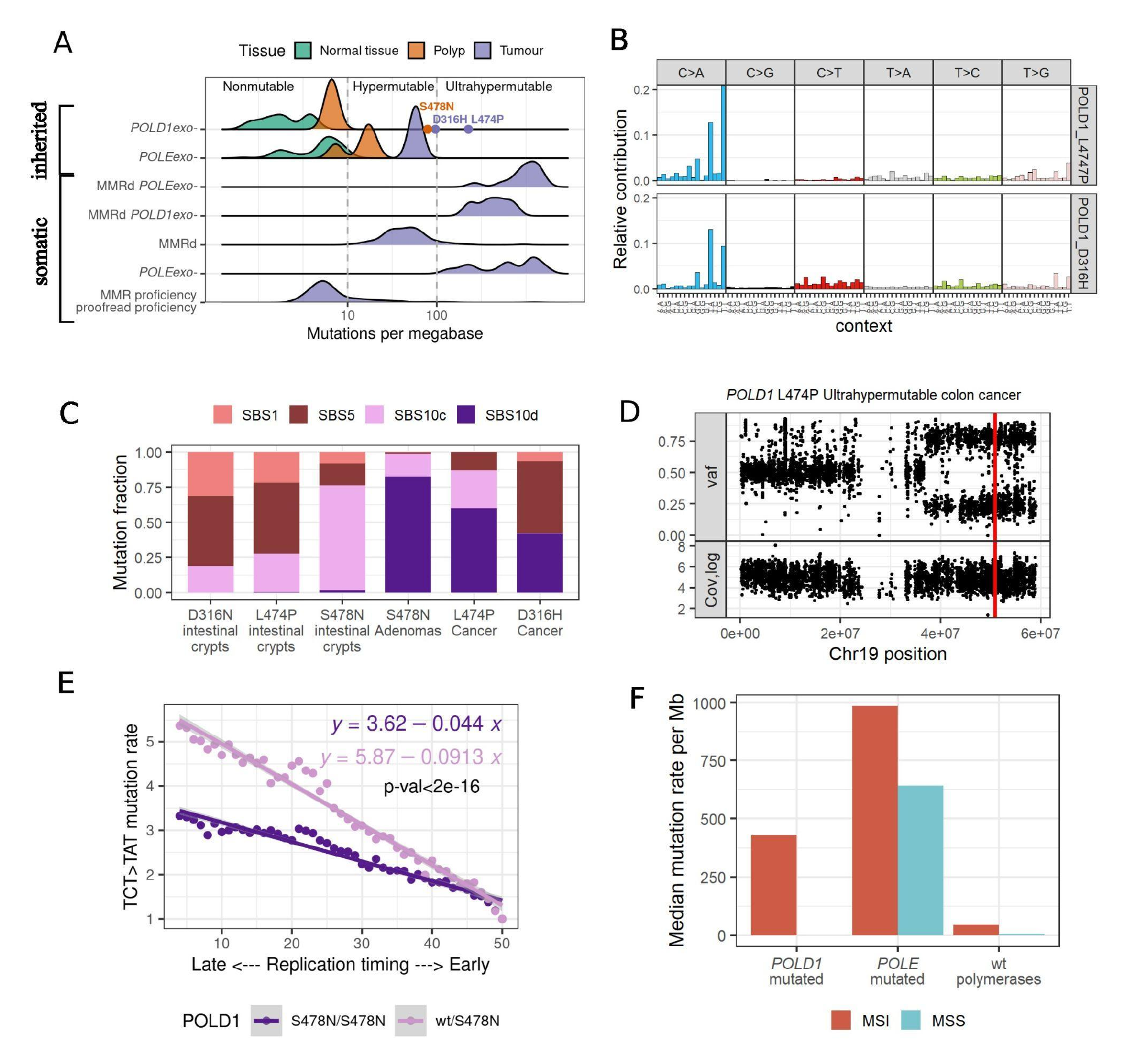
Bi-allelic inactivation of Polδ leads to hypermutability. **A**, Mutability of normal and cancer colon samples with inherited mutations in the exonuclease domain of *POLE* or *POLD1* and their formal classification based on the number of mutations per megabase (data from (Robinson et al. 2021)). Dots mark hypermutable and ultrahypermutable samples with inherited pathogenic mutations in *POLD1* (violet - two cancer samples sequenced in this study and orange – adenoma sample from (Robinson et al. 2021)). Mutation rates in uterine corpus endometrial carcinoma samples from The Cancer Genome Atlas (TCGA) with somatic inactivation of mismatch repair system (MMRd, MMR deficiency)) and/or proofreading activity of replicative polymerases (*POLE*exo-or *POLD1*exo-) are shown for comparison. **B**, 96-nucleotide context mutational spectrum of the ultrahypermutable cancer sample from the carrier of *POLD1* L474P and from an hypermutable cancer sample from a carrier of *POLD1* D316H. **C**, Fraction of mutations attributed to SBS10c and SBS10d mutational signatures in normal crypts, polyps (Robinson et al. 2021) and cancer samples from heterozygous carriers of *POLD1* variants. **D**, Variant allele frequency (top) and total coverage (bottom) along chromosome 19 in the tumor of a *POLD1* L474P carrier, showing cnLOH of the genomic region. The red line indicates the position of the *POLD1* L474P mutation. **E**, Fold change in mutation rate in bins of different replication timing compared to the bin with the earliest replication timing (RT). P-value corresponds to significance of interaction term between RT bin and homozygosity status in binomial regression model. **F**, Mutation rate in cancer exomes with heterozygotic somatic mutations in *POLE* or *POLD1* and varying status of the MMR system calculated for all available TCGA UCEC samples.

To explain this discrepancy, we considered the possibility of somatic inactivation of the exonuclease function in the second copy of *POLD1* in the hypermutable samples. Indeed, all three samples with elevated mutation rate, two tumors sequenced in this study and one polyp from (Robinson et al. 2021), had a copy neutral loss of heterozygosity (cnLOH) in the *POLD1* region. The cnLOH involves the loss of the wild-type *POLD1* allele, leading to homozygosity of the pathogenic *POLD1* variant in cells (Fig. 4D, Supplemental Fig. S10). Based on the overall number of structural variants identified in these three cases we estimated that the probability to observe cnLOH simultaneously in all three cases by chance is extremely low (p(cnLOH)=1.63*10^-8^, p(LOH) = 9.92*10^-7^) (Supplemental Table S4), suggesting the involvement of this rearrangement in thedevelopment of hypermutable phenotype. These observations suggest a high level of haplosufficiency of Polδ proofreading in human cells. Our findings show that the presence of a single copy of wild-type *POLD1* can prevent a strong increase in the mutation rate. In line with our results, haplosufficiency of *POLD1* was recently shown in yeast experiments (Zhou et al. 2021). A plausible explanation for this haplosufficiency is the ability of wild-type Polδ to proofread mismatches extrinsically, i.e., those produced by other mutant enzymes, thus preventing hypermutability. Our observations in normal and cancer tissues from heterozygous inherited *POLD1* variant carriers are concordant with observations in yeasts hinting that Polδ in human cells can also be able to proofread mismatches extrinsically. The abundance of TpCpA>A and TpCpT>A mutations in homozygous samples suggests that extrinsic Polδ proofreading is highly efficient for these types of mutations.

To identify and characterize the relative activity of the extrinsic proofreading effect of Polδ along the genome, we compared the relationship between replication timing and mutation rate in cells with homozygous and heterozygous *POLD1* L747P (our data) and S478N (Robinson et al. 2021). Both in hetero-and homozygous *POLD1*-mutated samples, mutation rates strongly depend on replication timing (Supplemental Fig. S11). However, in heterozygous samples, this dependence is more pronounced for both S478N and L474P (Fig. 4E, Supplemental Fig. S12), suggesting that the extrinsic proofreading effect of *POLD1* or its interaction with MMR is stronger in early replicating regions.

## DISCUSSION

Our findings indicate that inherited heterozygous mutations in the exonuclease domain of *POLD1* have a mild effect on somatic and germline mutation rates in humans, although *POLD1* exonuclease domain mutation carriers are predisposed to develop highly mutable cancers. Interestingly, we have observed that hypermutability in cancers or polyps is associated with loss of the wild-type *POLD1* allele. This result in human cells is in line with an extensive body of literature debating recessive effect of *POLD1* exonuclease deficiency in yeasts and in mice (Simon et al. 1991; Morrison et al. 1993; Zhou et al. 2021; Goldsby et al. 2002; Albertson et al. 2009).

It would be more intuitive to expect that *POLD1* exonuclease deficiency will have an additive effect on mutation rate, because mutated and non-mutated *Polδ* likely synthesize comparable amounts of DNA during replication of the lagging strand. Indeed, several potential explanations for the recessive behavior of *POLD1* mutations have been discussed in the literature, including a higher processivity of wild-type Polδ compared to the mutant allele (Simon et al. 1991), compensation by MMR (Goldsby et al. 2002), prevalent expression of the wild-type copy of *POLD1* (Daee et al. 2010) and others. A recent study in yeasts showed that the wild-type Polδ can correct mismatches produced by the mutant copy (extrinsic proofreading activity) (Zhou et al. 2021). Our data in human cells are consistent with this explanation, and we hypothesize that Polδ extrinsic proofreading helps prevent hypermutability in cells with heterozygous *POLD1* pathogenic variants (Fig. 5).

**Figure 5.**
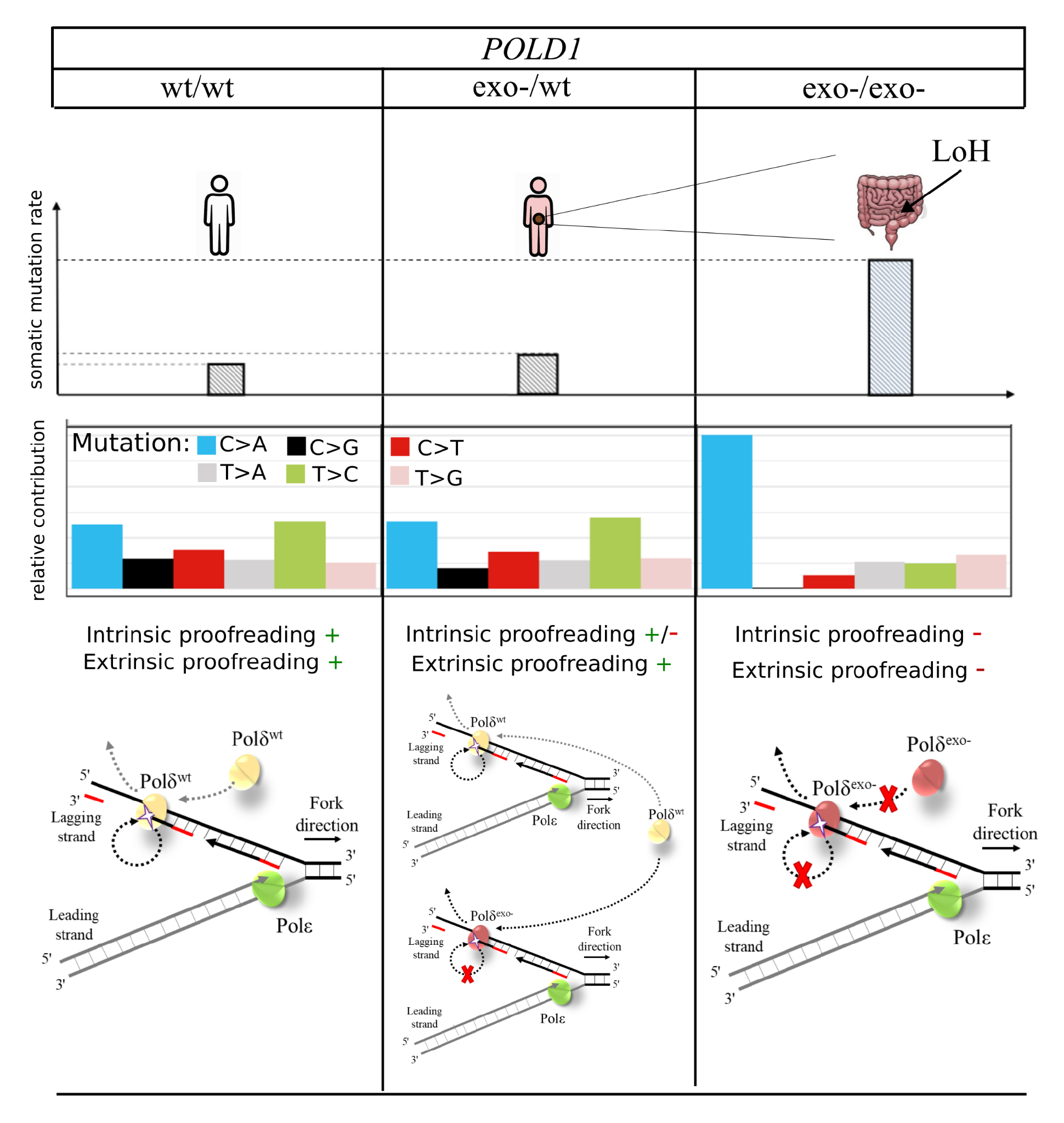
Schematic representation of the possible underlying mechanisms for monoallelic and biallelic inactivation of POLD1 proofreading activity.

Our results are well suited to explain the prevalence of different DNA repair deficiencies in cancer. Intuitively, a high mutation rate is a selected phenotype during cancer development. Here, we found that heterozygous mutations in the exonuclease domain of *POLD1* lead to a minor increase in mutation rate if other DNA repair systems are intact. Thus, heterozygous *POLD1* mutations should be almost neutral on a normal genetic background. Indeed, to our knowledge, there are no reported tumors where *POLD1* is somatically mutated but MMR is intact. Meanwhile, somatic inactivation of the second copy of *POLD1* on the background of a heterozygous inherited mutation is under strong positive selection and is likely a major avenue for cancer development. We could expect somatic MMR inactivation as an alternative mechanism; however, MMR deficiency (MMRd) is rarely found in cancers from individuals with inherited heterozygous *POLD1* mutations (Supplementary Table S5). This probably reflects the higher odds to mutate the second copy of *POLD1* compared to achieving MMRd.

Interestingly, tumors with somatic mutations in *POLD1* seem to be always accompanied by MMR deficiency (MMRd). In these tumors, analysis of signatures implies that MMRd precedes *POLD1* proofreading deficiency (Supplemental Fig. S13). MMRd alone increases the mutation rate, represents an advantageous genotype and could be fixed during tumor development; however, after a clonal expansion of MMRd, *POLD1* mutations will further increase the mutation rate by an order of magnitude (Fig. 4F) and thus should also be under positive selection. These two examples could explain why there are no tumors with only a heterozygous *POLD1* exonuclease mutation; either mutation of the second *POLD1* allele or an MMR deficiency (Haradhvala et al. 2018) are required for fixation of a somatic mutation affecting *Polδ* exonuclease domain.

Carriers of inherited pathogenic variants in the exonuclease domain of *POLD1* are mainly predisposed to colorectal and endometrial cancers, among other tumor types, which resembles the phenotypic features of the autosomal dominant hereditary cancer syndromes caused by *POLE* proofreading deficiency and MMRd. This tissue specificity may be explained by the fact that highly regenerative/proliferative tissues, such as the colon mucosa or the endometrial epithelium, are more sensitive to the deficiency in DNA repair mechanisms that deal with replication errors (polymerase proofreading defects, mismatches at the DNA replication forks, etc.), because inactivation of these systems will lead to extensive mutation accumulation (Sun et al. 2019). Another hypothesis for this tissue specificity might involve higher LOH levels in those epithelia. However, the fact that recessive polyposis and cancer syndromes caused by deficiency in other DNA repair mechanisms, e.g. those caused by inherited biallelic mutations in *NTHL1*, *MUTYH* and *MBD4*, also target the colorectal epithelium (and some predispose to endometrial cancer too), argues against it.

In summary, in this study we showed that heterozygous inherited *POLD1* L474P has a minor effect on mutation burden in germline and soma but leads to a prominent change in mutational spectra. Sequencing a large number of trios with mutations in genes involved in replication could help better understand the mutational mechanisms in the germline and identify the relative roles of replicative errors and DNA damage.

We also uncovered a recessive effect of polymerase delta proofreading deficiency in the context of cancer, suggesting that an inherited heterozygous *POLD1* exonuclease mutation may lead to cancer through a double hit mechanism, similarly to the somatic inactivation of the second MMR gene copy in Lynch syndrome patients (Porkka et al. 2017; Hemminki et al. 1994, 1; Sanchez de Abajo et al. 2006).

This and other findings suggest that the non-additive effect of polymerase proofreading and DNA repair genes inactivation on mutation rate is translated into recessivity and/or epistatic interactions in cancer.

## METHODS

### Study participants and Ethical approval

The family included in the study was recruited through the Hereditary Cancer Genetic Counseling and Molecular Genetics Lab at the University Hospital of Elche (Spain), where the clinical information and blood and skin punches were obtained. FFPE tumor material was obtained through the Valencian Biobank Network.

The study received the approval of the Ethics Committees of IDIBELL (PR235/16) and of the University Hospital of Elche (PI37/2018).

### Experimental procedure to assess somatic mutation accumulation in fibroblasts

#### Fibroblast obtention and immortalization

A skin punch biopsy was obtained from individuals III.2, III.4, IV.1, IV.2, IV.3, IV.4, IV.5, and IV.6 (Figure 1; Supplementary Table 1). The obtained biopsies were kept on culture media [DMEM with 10% fetal bovine serum and 100U/ml penicillin-streptomycin (Gibco, Thermo Fisher Scientific, Walthman, MA] at 4°C until processing. After a 1X PBS wash, the sample was incubated ON at 37°C with digestion media (DMEM supplemented with 160 U/ml collagenase and 1.25 U/ml dispase) and then disaggregated by pipetting. The sample was washed with culture media and seeded in one well of a 12-multiwell plate and maintained at 37°C in a 5% CO_2_ atmosphere. Cells were expanded for two passages before immortalization with lentiviral transduction with hTERT.

The lentiviral plasmid carrying the catalytic subunit of human telomerase protein (pLVX-IRES-hTERT-tdTOMATO) as well as envelope and packaging plasmids (pPAX2 and pMD2G) were kindly provided by Dr. Manel Esteller. Lentiviruses were produced transfecting HEK293 cells growing on a T75 flask with 20ug of total DNA and Lipofectamine 2000 (Thermo Fisher Scientific) for 16h. The culture medium was changed, collected at 72h and filtered using a 0.45 µm filter. The lentivirus enriched-medium was immediately used to transduce the human fibroblasts growing on a T25 flask in the presence of 8µg/ml polybrene (Sigma-Aldrich, San Luis, MO). Infected fibroblasts were maintained in culture until having enough cells for cell sorting. Cells expressing hTERT-tdTOMATO were enriched by fluorescent activated cell sorting (FACS). Cells were analyzed with the cell sorter MoFlo Astrios (Beckamn Coulter, Brea, CA) using a 561nm laser. Immortalized fibroblasts were maintained in culture media at 37°C in a 5% CO_2_ atmosphere and split at 1:3 ratio.

#### Single cell isolation and clonal expansion

To generate single-cell clones, fibroblasts enriched in hTERT-tdTOMATO were single-cell sorted using the MoFlo Astrios and the 561 nm laser. Each single cell was automatically plated in a well of a 96-well plate in the presence of 200 µl of culture medium. The individual cultures were followed up to ensure clonal expansion. Each pool of cells was passed to growing sizes of culture plates up to T25 flasks. The first confluent T25 flask was considered the starting passage (p0). Two clones per individual were maintained in culture for 30-45 additional passages (Supplementary table 2) as described above. One clone per individual was sequenced.

### DNA extractions

Peripheral blood DNA was extracted using the FlexiGene DNA kit (Qiagen, Valencia, CA). DNA from fibroblasts was obtained with the Quick-DNA Miniprep Plus kit (Zymo Research, Orange, CA). DNA from buccal swabs obtained with Isohelix swab packs and maintained in BuccalFix tubes (Isohelix, Cell Projects Ltd, UK) was extracted using a standard phenol-chloroform protocol. DNA from formalin-fixed paraffin-embedded (FFPE) tumor samples was isolated with the kit QIAamp DNA FFPE tissue Kit (Qiagen). All extractions were carried out following the manufacturers’ instructions.

### Whole-genome sequencing (WGS)

DNA preparation for WGS was performed with the TruSeq Nano DNA Library, and sequencing was carried out in a NovaSeq 6000 150 PE (2×150 bp). Sequencing was performed at a minimum coverage of 90 Gb (30x) for the germline studies in the family members, and at a minimum coverage of 150 Gb (50x) for the experiments with fibroblasts (P0 – ∼P40 passages of the fibroblasts’ cultures). Sequencing was performed at Macrogen (Macrogen Inc, Seoul, South Korea).

### Whole-exome sequencing

Exome sequencing was performed in FFPE tumor DNA of individual IV.6. Exome capture was performed with Kappa HyperExome Probes (Roche) and sequenced in a NovaSeq 6000 S1 (2×100bp). Sequencing was performed at Centro Nacional de Análisis Genómico (CNAG, Barcelona, Spain).

### Analysis of MMR status in tumors

MMR status in tumor tissue was assessed by standard methods, using immunohistochemistry of MMR proteins MLH1, MSH2, MSH6 or PMS2, and/or by analysis of microsatellite instability (MSI) by PCR-based analysis of microsatellite markers.

### Variant calling in fibroblasts and post-processing filters

Sequenced reads were aligned to hg19 reference human genome downloaded from UCSC (https://hgdownload.soe.ucsc.edu/downloads.html#human) using Burrows–Wheeler alignment (BWA-MEM) (Li 2013).

Somatic mutations in single-cell derived colonies were called using Mutect2. DNA genome sequencing data available from normal tissue (blood, buccal swab or fibroblasts) from the corresponding individual was used as matched normal DNA. Panel of normals (--panel-of-normals Mutect2 argument) was created from all available sequenced blood samples. The population allele frequencies in gnomAD were used as a prior for germline variant detection (-- germline-resource Mutect2 argument). Additional filter for the variant allele frequency observed in sequenced data was applied: only mutations with variant allele frequencies (VAFs) between 0.25 and 0.75 were selected for the analysis.

Somatic mutations accumulated during the experiment were called using Mutect2. Mutations present in the end point (P40) but absent in the start (P0) were selected. To filter out recurrent technical artifacts, all available sequencing data from samples that did not correspond to the mutation accumulation experiment were used to create a panel of normals (--panel-of-normals Mutect2 argument). The population allele frequencies in gnomAD were used as a prior for germline variant detection (--germline-resource Mutect2 argument).

### Mutational spectra and PCA analysis

The 96-mutational spectrum was calculated dividing the number of mutations of a particular type in fixed 3-nucleotide contexts by the total number of mutations of that type. Principal component analysis (PCA) was used on these 96-dimensional vectors for mutations obtained from fibroblast colonies. Mutational spectra obtained by the same procedure across different datasets (Fig. 2f, Fig. 3c,e, Fig.3g,h) were projected onto the obtained PC space.

### Extraction of *de novo* signatures

Mutational signatures in fibroblasts were extracted *de novo* and then decomposed to COSMIC signatures using SigProfilerExtractor (Islam et al. 2020). The number of mutations attributed to each COSMIC signature was obtained as output.

For mutations accumulated in fibroblast colonies during the experiment, SigProfilerExtractor was run with the following parameters: minimum_signatures=1, maximum_signatures=15, nmf_replicates=300. The solution with the most stable signatures (n_signatures = 2) was selected. Two de novo extracted signatures were decomposed in 6 reference COSMIC signatures. The initial mutational pattern of each sample was refitted to COSMIC signatures using the SigFit package. To avoid overfitting the subset of signatures for decomposition, the analysis was limited to the six signatures predicted by SigProfilerExtractor: SBS5, SBS10c, SBS36, SBS37, SBS45 and SBS93.

The ‘sigfit’ R package (Gori and Baez-Ortega 2018) was used to estimate exposure to mutation signatures in intestinal crypts, adenomas and cancer samples with inherited *POLD1* mutations. In this case, the signatures fitted were limited to SBS1, SBS5, SBS10c and SBS10d.

The ‘mSigAct’ package was used to estimate the significance of SBS10c presence (Ng et al. 2017).

### Calling of germline *de novo* mutations and post-processing filters

Sequenced reads were aligned to the hg19 reference human genome downloaded from UCSC using Burrows–Wheeler alignment (BWA-MEM). *De novo* mutations were called using standard GATK4 best practices pipeline. All samples were jointly called, but only high-confidence mendelian violation sites with minimum GQ=20 for each trio member were selected. Subsequently, additional filters were applied: i) coverage of each trio member ≥10; ii) absence of reads confirming alternative allele in parents; iii) number of reads confirming reference and alternative allele in proband ≥5; iv) allele frequency of the alternative allele in the proband ≥0.3. Additionally, clustered mutations (distance between mutations ≤1000bp) were filtered out as potential false positives. Mutations present in any other family member were also excluded. To eliminate false positive mutations coming from sequencing of proband fibroblasts, we used other fibroblasts from the same individual (used in MA experiments) as additional confirmation of mutation presence: true *de novo* mutations have to be present in all tissues of the proband, including the other fibroblast sample. This additional filter mainly removed variants from the left tail of the variant allele frequency distribution of potential *de novo* mutations, thus keeping the distribution more symmetric around 0.5 (Supplemental Fig. S14). To evaluate the performance of the filter, we took the data from our preliminary sequencing of blood samples for two trios from the family (IV.5 and IV.6 probands) and compared the *de novo* mutations obtained in the two attempts. We found out that for the IV.6 offspring, this additional filter removed 1257 out of 1360 variants. Only 6 of them had been proposed as *de novo* mutations in the first round of trio sequencing. For the IV.5 individual, the filter removed 56 out of 121 candidates and none of them had been called as *de novo* mutations in the preliminary sequencing (Supplemental Fig. S15). Blood samples for each individual in the trios had been preliminary sequenced with 20X coverage, *de novo* mutations had been called using PhaseByTransmission GATK3 tool (--prior = 1e-4) and sites with violation from mendelian inheritance were selected.

### Enrichment of *de novo* mutations in *POLD1* contexts

According to analysis of mutations in fibroblasts, four contexts were enriched in mutations in carriers of *POLD1* L474P variant: Cp<u>C</u>pT>A, Tp<u>C</u>pT>A, Ap<u>T</u>pT>A and Cp<u>T</u>pT>G.

The calculated proportion of mutations in these contexts in *de novo* mutations of published trios equalled 2.4%. This is thus the expected proportion for such mutations in offspring of wild-type *POLD1* parents. We then calculated the expected number of such mutations in each offspring in our experiment by multiplying the expected proportion by the total number of observed mutations in offspring. The sum of expected numbers calculated by all offspring of father-carriers of *POLD1* L474P variant was compared to the observed sum to estimate the enrichment. Rate-ratio test was used for comparison of observed vs. expected data.

The proportion of mutations in *POLD1* contexts in somatic cells was calculated using mutations from *POLD1* L474P fibroblasts excluding IV.6 and IV.1 samples.

### Simulation of *de novo* mutations and test for the presence of mutators in the population

To test how many offspring of wild-type parents in the sample have –PC1 values higher than the mean value of –PC1 in the offspring of fathers harboring *POLD1* L474P by chance, we generated an artificial dataset of trios by randomly sampling *de novo* mutations according to their fractions in homogeneous underlying spectra. To obtain the underlying spectra, we used data from 6233 offspring from two publicly available datasets, aggregated all *de novo* mutations together and calculated the proportion of each mutation type in each possible 3-nucleotide context. For each offspring in the dataset, we counted the observed number of *de novo* mutations, and sampled the same number of mutations from the obtained spectrum. For each simulated set of *de novo* mutations, we created the vector of mutational probabilities and estimated the value of the -PC1 component by projecting in the PC space obtained previously from the analysis of fibroblasts. We repeated the same procedure generating two or more offspring for each family and averaging the –PC1 among siblings.

Similarly, a sample of trios with a mixture of offspring of *POLD1* L474P carriers was simulated. For 95% of the offspring we sampled mutations according to their probability in the wild-type trios and for the remaining 5% of samples, we sampled mutations based on their probability of occurring in the offspring of *POLD1* L474P fathers in the studied family. We then calculated the mutational spectrum per simulated individual and the -PC1 values.

For other mutagenic processes (Polε proofreading inactivation, defective base excision repair) *de novo* mutations for mutators (proportion of mutators varied from 0 to 10%) were sampled as a mixture of mutations from wild-type trios spectrum and spectrum of the corresponding known COSMIC signature attributed to the mutagenic process (SBS10a, SBS36 correspondingly). Cosine similarity with the COSMIC signature was used to estimate the presence of this signature in the simulated spectrum of the sample. Kolmogorov-Smirnov test was used to estimate the significance of difference between simulated subsets: subset of wt trios vs subset with mutators (Supplemental Fig. S9).

For simulation in Supplemental Figure S7 we created a synthetic dataset of *de novo* mutations for 5000 wild type trios as a background. The number of mutations per individual was generated from negative binomial distribution (mu=60, size=80) and mutations were sampled from the observed spectrum of *de novo* mutations in wt trios. For 5 samples in addition to this procedure we added mutations sampled from SBS10c signature spectrum. The number of additional mutations was generated from the Poisson distribution with lambda equal to 0.15 multiplied by the number of mutations from the wt spectrum. Projection to PC1 and proportion of *POLD1*- specific contexts were used to estimate presence of mutations introduced by additional mutagenic process (Polδ proofreading inactivation).

### Calling of mutations in tumors from *POLD1* pathogenic variant carriers

Sequenced reads were aligned to the hg19 reference human genome downloaded from UCSC (https://hgdownload.soe.ucsc.edu/downloads.html#human) using Burrows-Wheeler alignment (BWA-MEM). Somatic mutations in tumor samples were called against normal samples from the same individual using standard GATK4 best practices pipeline. The Learn Orientation Bias Artifacts tool was used to control for orientation bias, which is critically important for FFPE samples. Mutations with variant allele frequency <0.15 were filtered out, as low-frequency variants are known to be enriched in formalin fixation artifacts in FFPE samples (Bhagwate et al. 2019).

### Epigenetic covariates for the extrinsic proofreading effect of Polδ

Genome was splitted in 100-kb non-overlapping windows. The number of mutations and target sites was calculated in each window. The replication timing (RT) is highly conserved between human tissues and cell types (Ryba et al. 2010; Pope et al. 2014). Here we used the mean replication timing for each window obtained using Wavelet-smoothed Signal for Repli-seq data for HeLa-S3 cell line (GSE34399). Windows were splitted in bins according to RT mean value; mutation rate in each bin was calculated as a sum of all mutations in windows in this bin divided by the sum of target sites. The obtained mutation rate in each bin was normalized for the mutation rate in the bin of the earliest replication time.

For S478N samples whole genome data was available. For heterozygous S478N, we used data from 32 samples from 6 individuals (48052 mutations in total). For homozygous S478N we were able to use only 1 sample from 1 individual with a total of 66774 mutations. Mutational data were obtained from a previously published dataset (Robinson et al. 2021). Only Tp<u>C</u>pT>A/ApGpA>T mutations were analyzed and 50 RT bins were used. To estimate statistical significance of the difference, we ran binomial regression with interaction using RT bin and homozygosity of S478N variant as predictors. The p-value for the interaction term is provided in Fig 4E.

For L474P samples whole exome data was available, all mutations were analyzed together and RT was splitted in 3 bins only.

### LOH rate estimation

We called LOH regions in all three samples independently using CNVkit (Talevich et al. 2016) and calculated the rate of LOH in each sample dividing the number of sites with LOH by the total number of called target sites. To get an unbiased estimation, we excluded chr19 from this estimation as we wanted to check the independence of LOH in the *POLD1* gene located in that chromosome. The probability to observe LOH in all three samples was obtained as the product of three probabilities.

## DATA ACCESS AND SOFTWARE AVAILABILITY

DNA sequencing data are deposited in the European Genome-Phenome Archive (EGA) with accession code EGAS00001006434. Called de novo and somatic mutations are available online (https://figshare.com/account/home#/projects/156404).

Code required to reproduce the analyses in this paper is available online (https://github.com/andrianovam/POLD1-project_scripts).

## ACKNOWLEDGMENTS

We would like to thank the members of the family under study for their selfless participation, and Dr. Anouk Jaen Larrieu of the Dermatology Department of Elche University Hospital, who performed the skin biopsies for the obtention of the fibroblasts.

V.B.S., F.A.K., A.S.K., G.A.B. and L.V. conceived the project and designed the experiments. M.A.A. and V.B.S. conducted bioinformatic and statistical analyses. M.T., P.M., and G.A. conducted the experiments in the fibroblasts from family members (immortalization, cell culture and single cell isolation) under L.V.’s supervision, and performed DNA extractions from blood, buccal swabs and tumor samples. A.B.S-H. and J.L.S. obtained the clinical information, informed consents and samples from the family under study. M.A.A., V.B.S. and L.V. wrote the original manuscript. F.A.K., A.S.K. and G.A.B. consulted on the project and edited the manuscript.

