## Supplementary Material for "Extended family with an inherited pathogenic variant in polymerase delta provides strong evidence for recessive effect of proofreading deficiency in human cells"

### Table of contents

Supplemental Note.

Supplemental Figure S1.

Supplemental Figure S2.

Supplemental Figure S3.

Supplemental Figure S4.

Supplemental Figure S5.

Supplemental Figure S6.

Supplemental Figure S7.

Supplemental Figure S8.

Supplemental Figure S9.

Supplemental Figure S10.

Supplemental Figure S11.

Supplemental Figure S12.

Supplemental Figure S13.

Supplemental Figure S14.

Supplemental Figure S15.

Supplemental Table S1.

Supplemental Table S2.

Supplemental Table S3.

Supplemental References

**Supplemental Note.**

Evidence that supports the pathogenicity of *POLD1* L474P and its influence on the proofreading activity of Polδ. The ACMG/AMP guidelines adapted to the *POLD1* gene (Richards et al. 2015; Mur et al. 2020) classifies L474P as likely pathogenic (evidence and rule codes highlighted in blue, between brackets).

- Germline carriers show the phenotypic characteristics of the hereditary cancer syndrome associated with polymerase proofreading deficiency (PPAP). The variant segregates with the cancer and/or polyposis phenotypes (Valle et al. 2014; Bellido et al. 2016; Ferrer-Avargues et al. 2017; Palles et al. 2022). [PP1_strong: when co-segregation is observed in ≥7 meioses in ≥2 families (reported L474P families: 9 meiosis in 4 families)]
- Absence in population databases (zero individuals in gnomAD v.2 containing 125,748 exomes and 15,708 genomes, or in gnomAD v.3 containing 76,156 genomes) [PM2_supporting: L474P is absent in control populations]
- It affects a highly conserved residue within the Exo IV motif of the exonuclease domain of Polδ. The variant is predicted pathogenic by computational tools ( REVEL (metapredictor) score = 0.913; "1" being the highest possible value for pathogenicity. [PP3: Missense ED variants with a REVEL score ≥0.5]
- Modification of the L474 residue leads to proofreading dysfunction (experiment performed in yeast (*Saccharomyces cerevisiae*) (Murphy et al. 2006)). Own experimental data in *Schizosaccharomyces Pombe* show an effect of L474P on the proofreading activity (hypermutator phenotype) (Supplemental Fig. S1)). [PS3_supporting: hypermutator phenotype observed in a yeast-based experimental system]

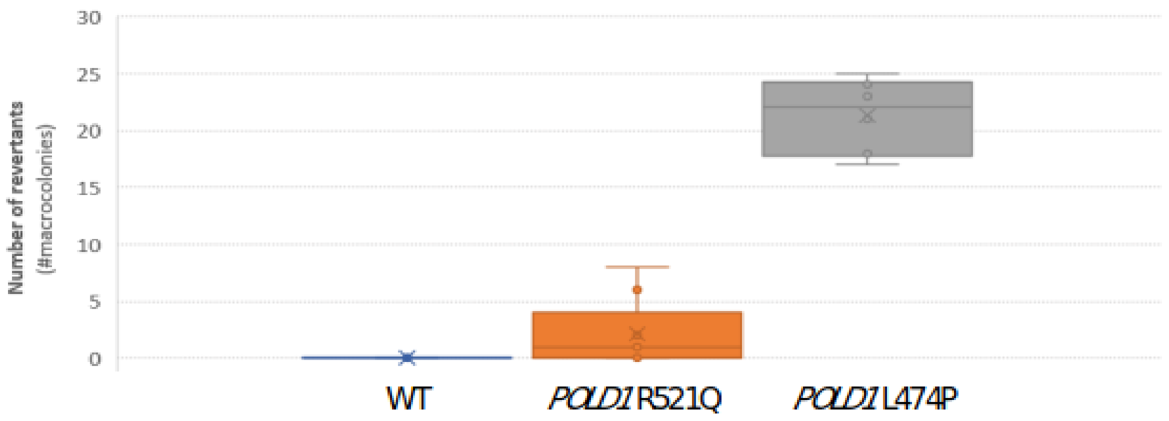


**Supplemental Figure S1.**

Number of revertant colonies (per plate) in different genotype backgrounds (wild-type, *POLD1* R521Q and *POLD1* L474P) in *Schizosaccharomyces pombe*. Colony growth occurs when spontaneous mutations cause the reversion of the ade6-485 allele in the corresponding yeast strain due to polymerase proofreading deficiency. No growth of revertant colonies is expected in the WT strain. The mean and standard deviation was obtained from two independent experiments performed in triplicate. The three variants were assayed in parallel. The methodology used was described previously (Mur et al. 2020). *POLD1* variant nomenclature in the figure corresponds to the human gene. R521Q is a variant of unknown significance in the exonuclease domain of *POLD1*.

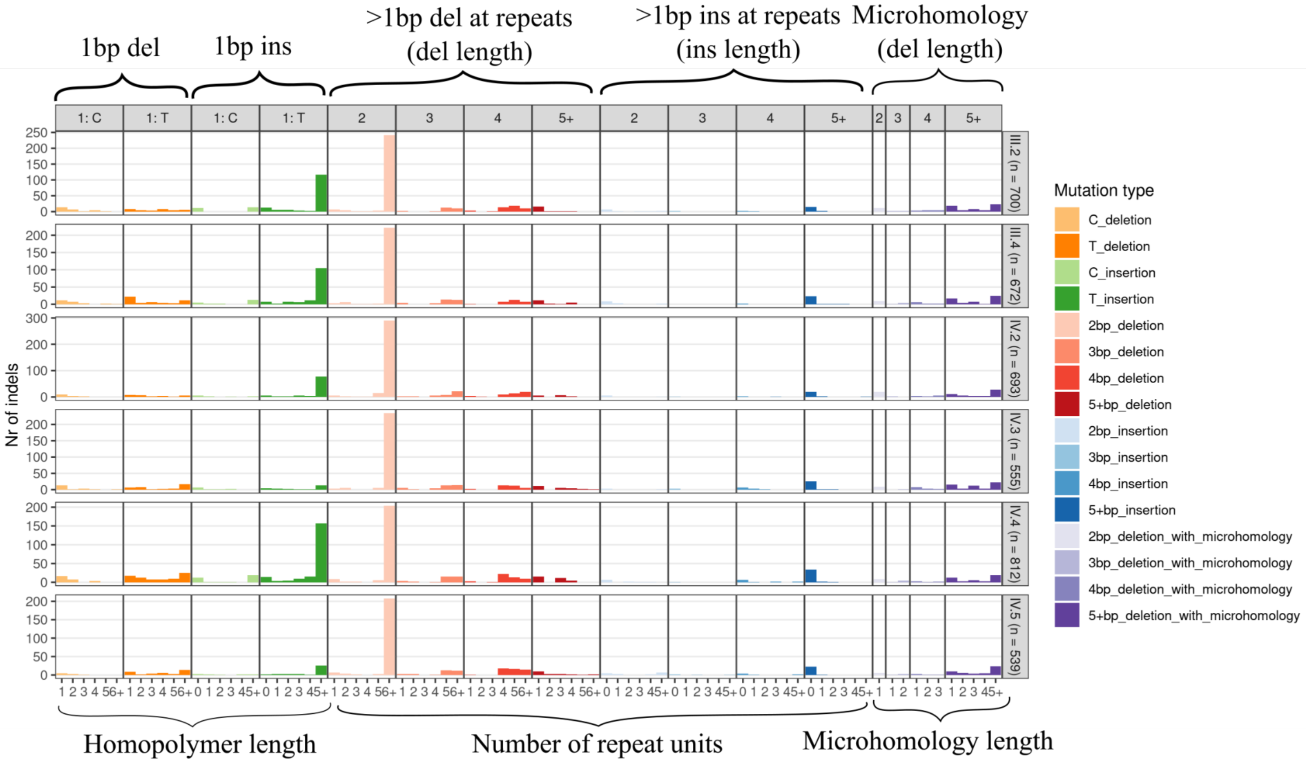


#### Supplemental Figure S2.

Insertions and deletions accumulated during the experiment in fibroblasts colonies with heterozygous *POLD1* L474P. Values in brackets near the sample names correspond to overall observed number of insertions and deletion.

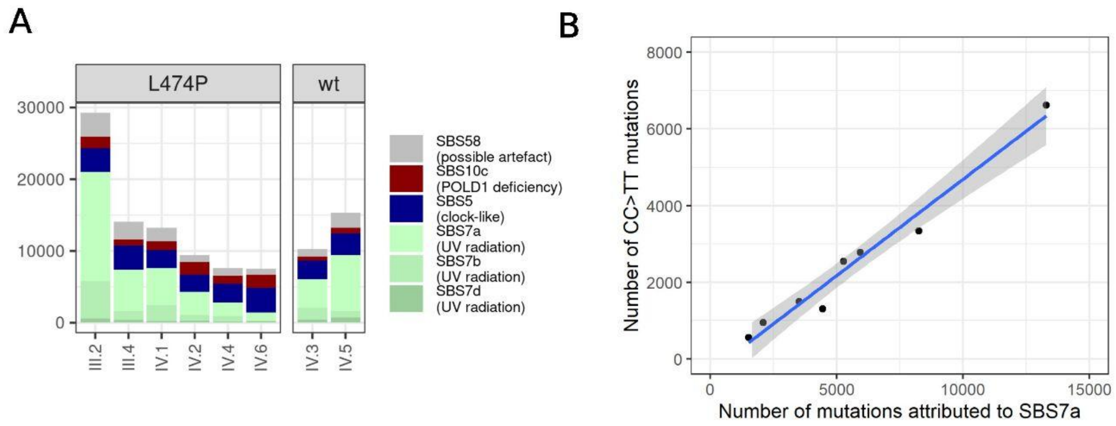


**Supplemental Figure S3.**

Mutations accumulated in skin fibroblasts during life. **A,** COSMIC mutational signatures found in single-cell colonies from skin fibroblasts. The list of signatures was obtained using de novo extraction by SigProfilerExtractor and subsequent refit using SigFit. **B,** Correlation of the number of mutations attributed to SBS7a and number of CC>TT double substitutions in skin fibroblasts colonies.

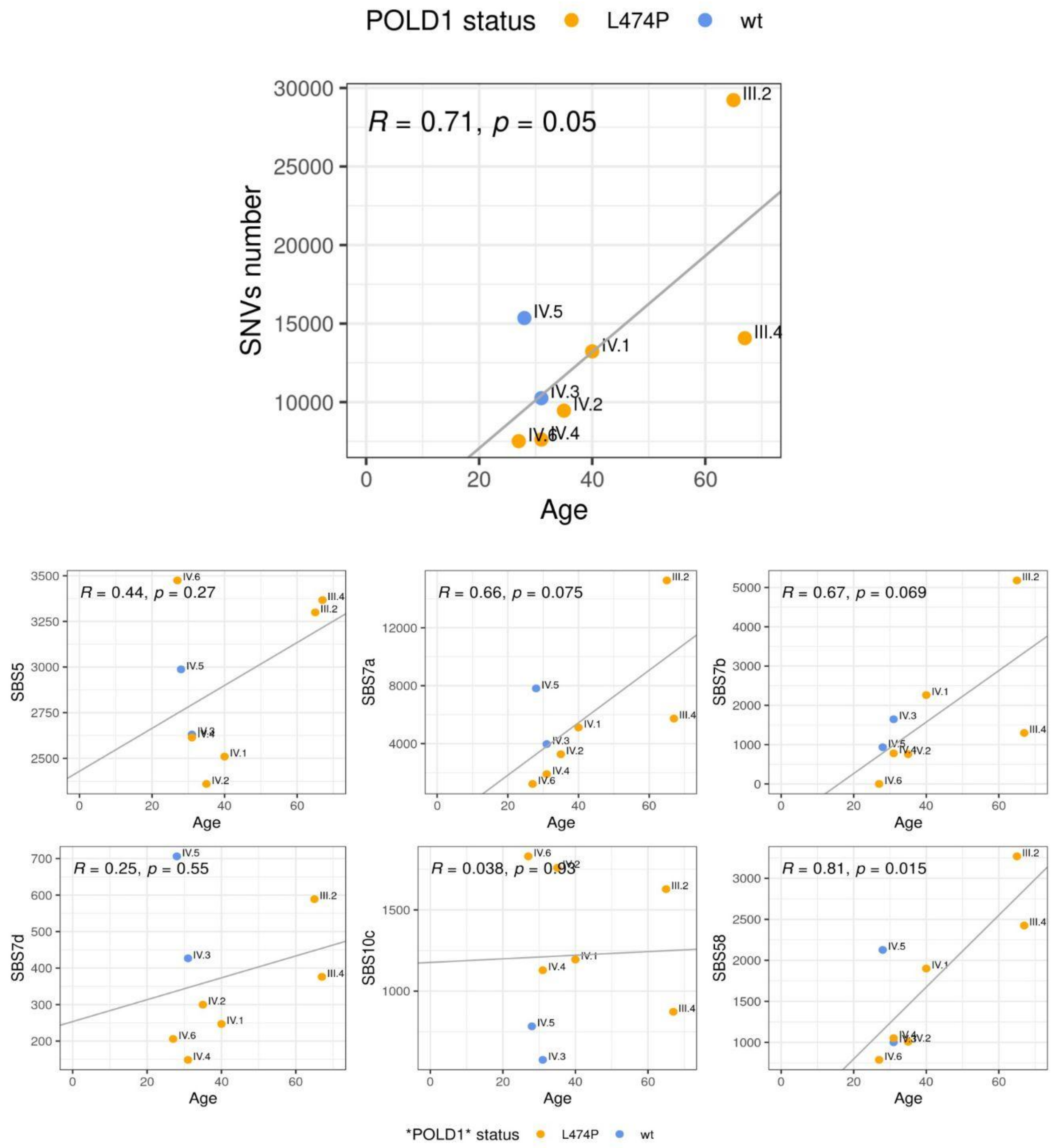


**Supplemental Figure S4.**

Correlation of the number of mutations with individual age at the time of biopsy.

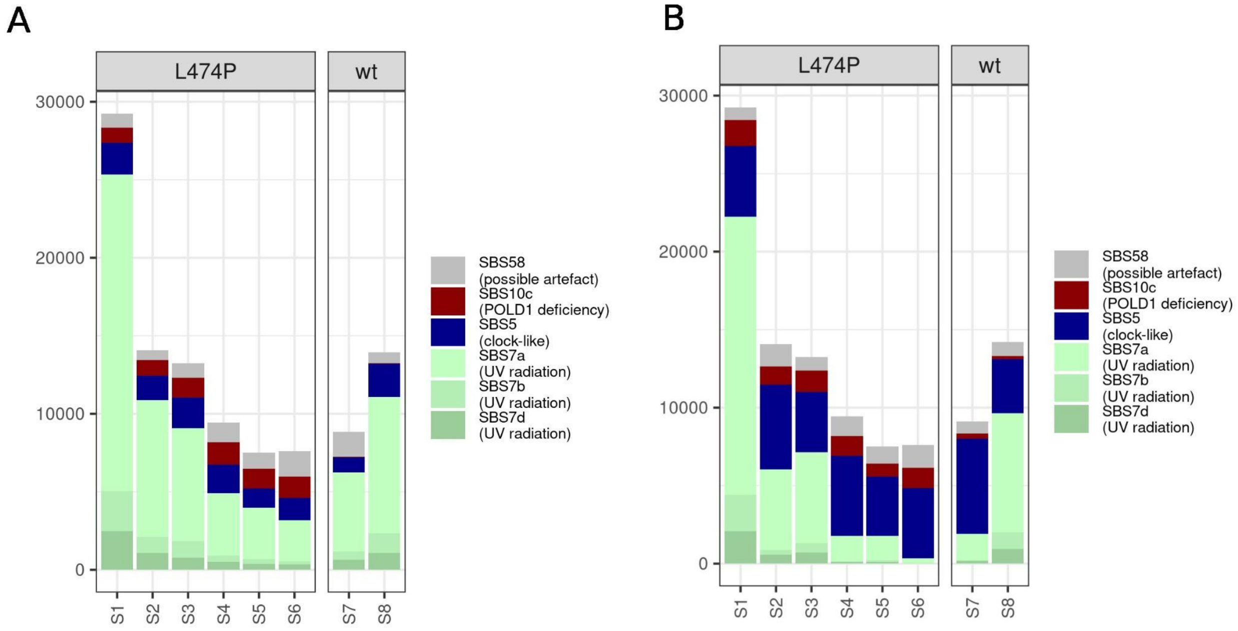


#### Supplemental Figure S5.

Signatures observed in synthetic samples after refitting procedure by SigFit. Synthetic samples were generated with the number of mutations corresponding to real samples. Wild type samples were generated using SBS5 (500 mutations), SBS58 (1000 mutations) and SBS7a,b,d (other mutations). In L474P samples 1000 mutations from SBS10c were added. **A,** 1000 random mutations representing noise were added, **B,** 4000 random mutations were added.

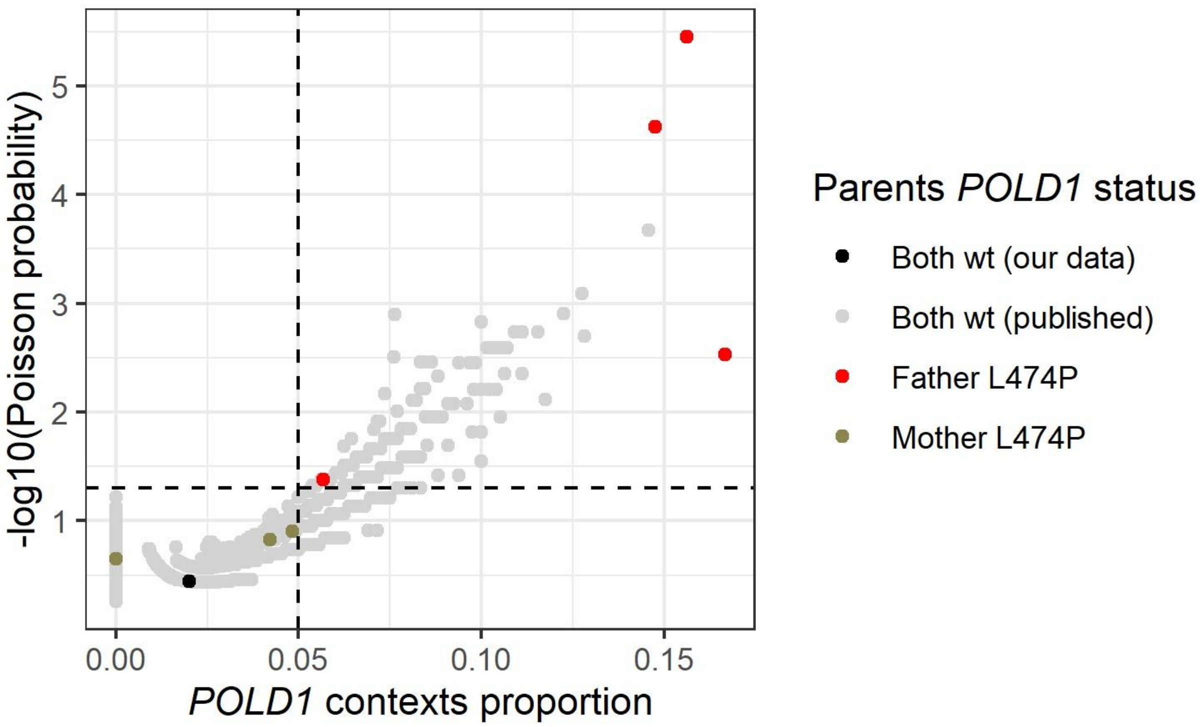


**Supplemental Figure S6.**

Proportion of mutations in four 3-nucleotide contexts specific for mutated *POLD1* against Poisson probability of observed number of mutations in these contexts. Dashed horizontal line marks p-value=0.05. Dashed vertical line marks 90%-percentile of contexts proportion in published trios (Halldorsson et al. 2019; An et al. 2018).

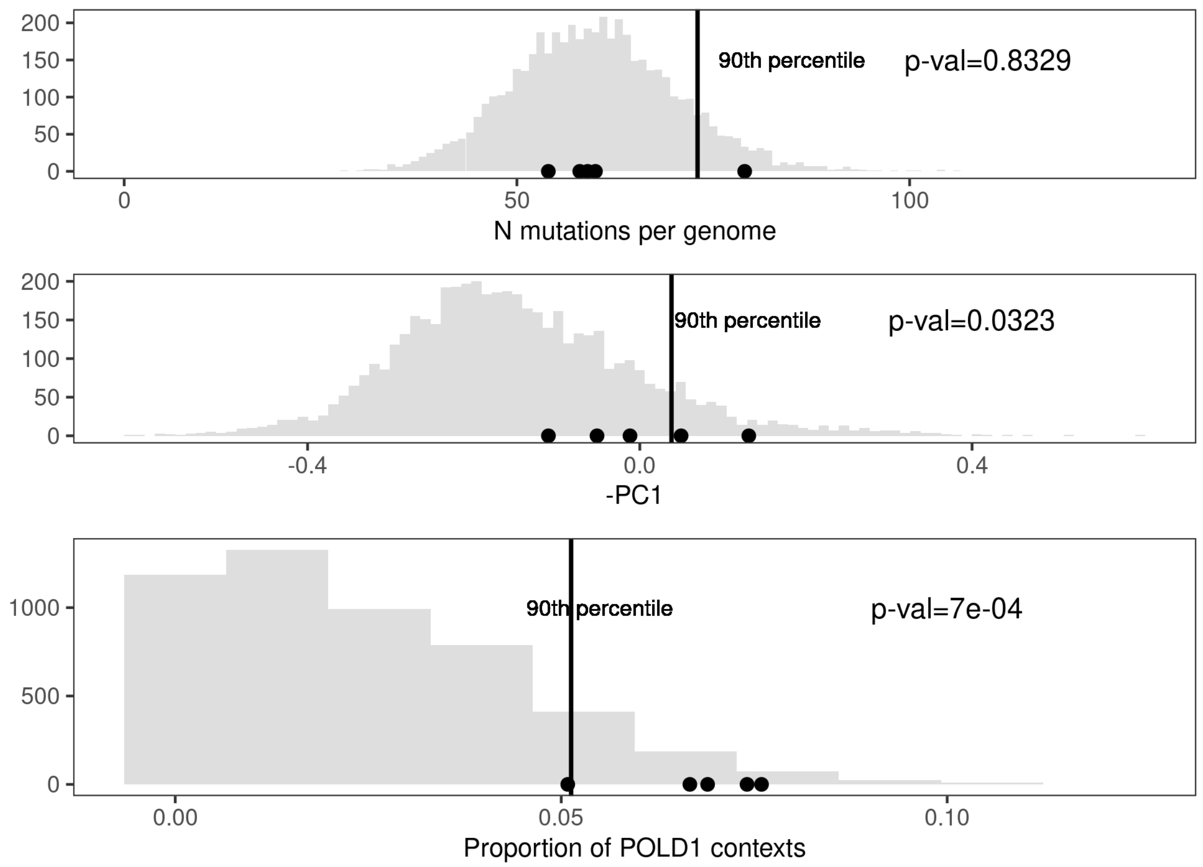


**Supplemental Figure S7.**

Representation of the number of mutations (top), -PC1 values (middle), and proportion of *POLD1*-specific contexts (bottom) in synthetic dataset of wt trios (gray distribution) and trios with 15% of mutations added from spectrum of SBS10c COSMIC signature. The p-value for Kolmogorov-Smirnov test is shown.

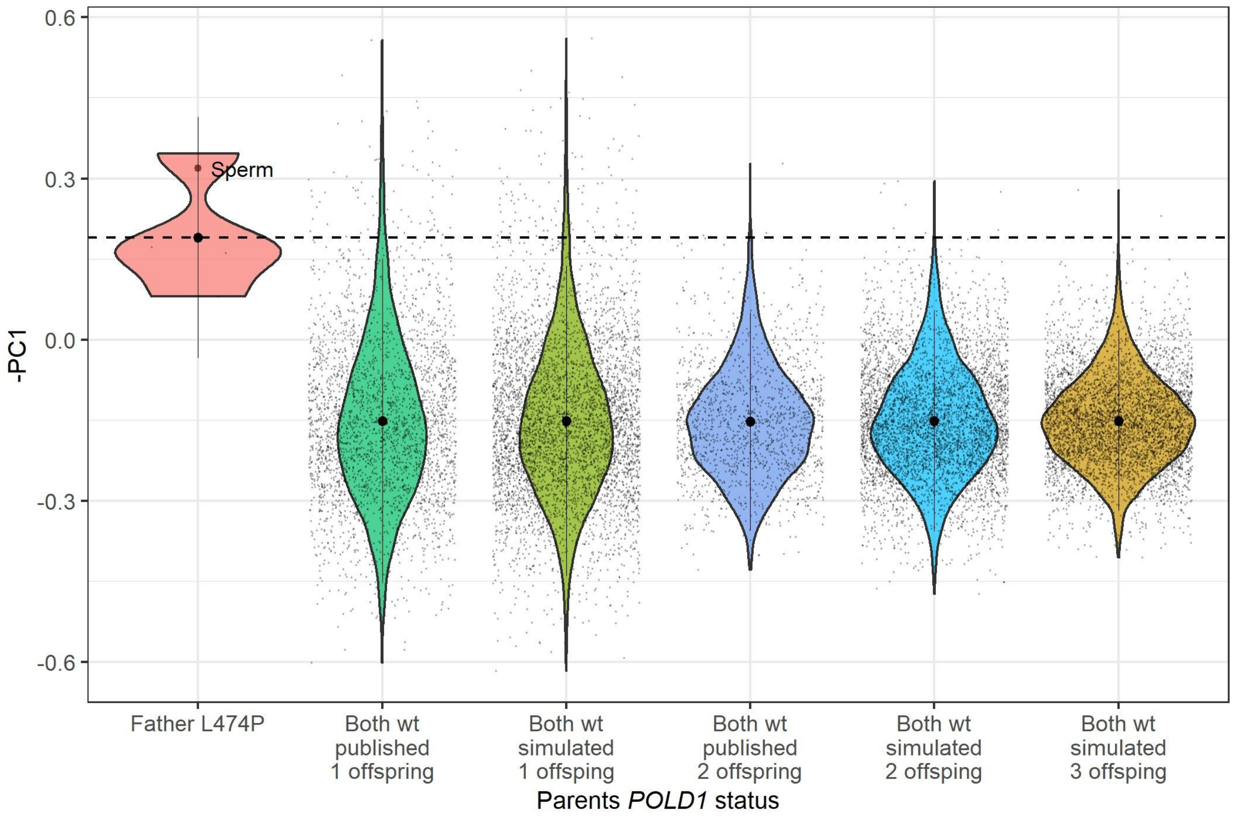


**Supplemental Figure S8.**

Proportion of POLD1-associated contexts in de novo mutations in trios with *POLD1* L474P carrier father and in observed and simulated families with different numbers of offspring. For families with more than one offspring – PC1 value averaged among offspring is shown for each family. Dashed line corresponds to mean –PC1 value in the offspring of fathers harboring *POLD1* L474P.

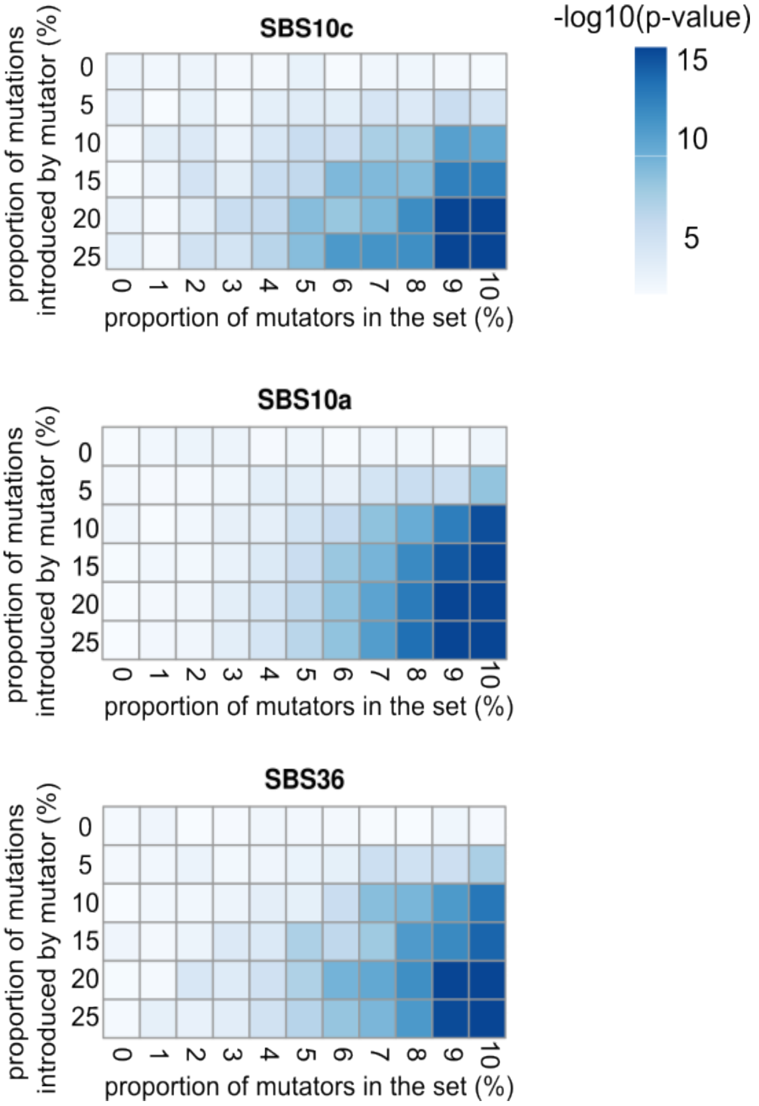


**Supplemental Figure S9.**

Statistical significance (two-sample Kolmogorov-Smirnov test, -log10 p-value) of difference in distributions of cosine similarities to target mutational signature (SBS10c, SBS10a or SBS36) between synthetic wild type population and synthetic population with presence of mutators. The proportion of mutators in the synthetic population varies between 0% and 10% (x-axis) and the proportion of mutations contributed by additional mutagenic process in the mutators varies between 0% and 25% (y-axis).

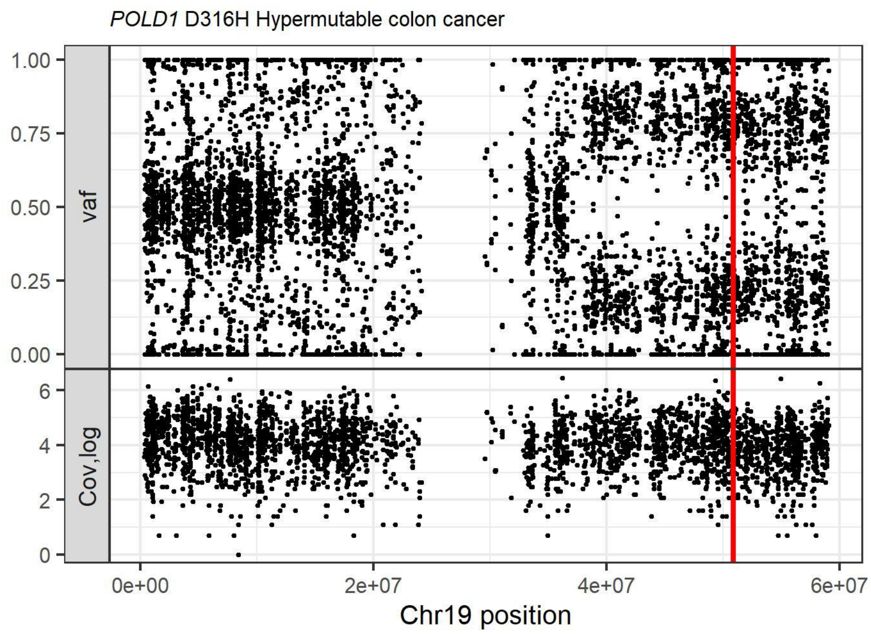


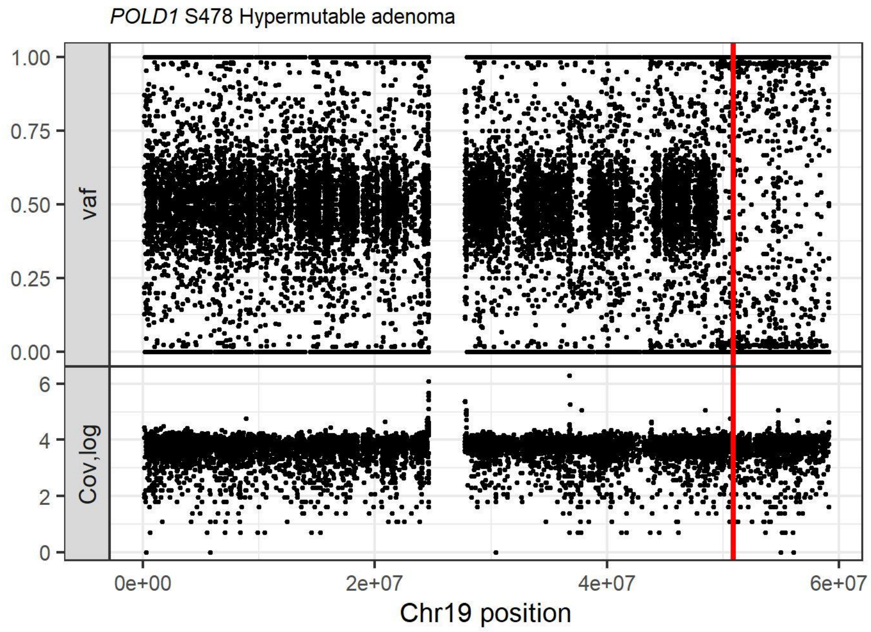


#### Supplemental Figure S10.

cnLOH in tumor from germline carrier of *POLD1* D316H and in a hypermutable polyp from a germline carrier of *POLD1* S478N (Robinson et al. 2021). The top plot for each sample shows variant allele frequency of germline variants, the bottom plot shows the logarithm of coverage for each variant. The red line indicates the position of the inherited *POLD1* pathogenic mutation.

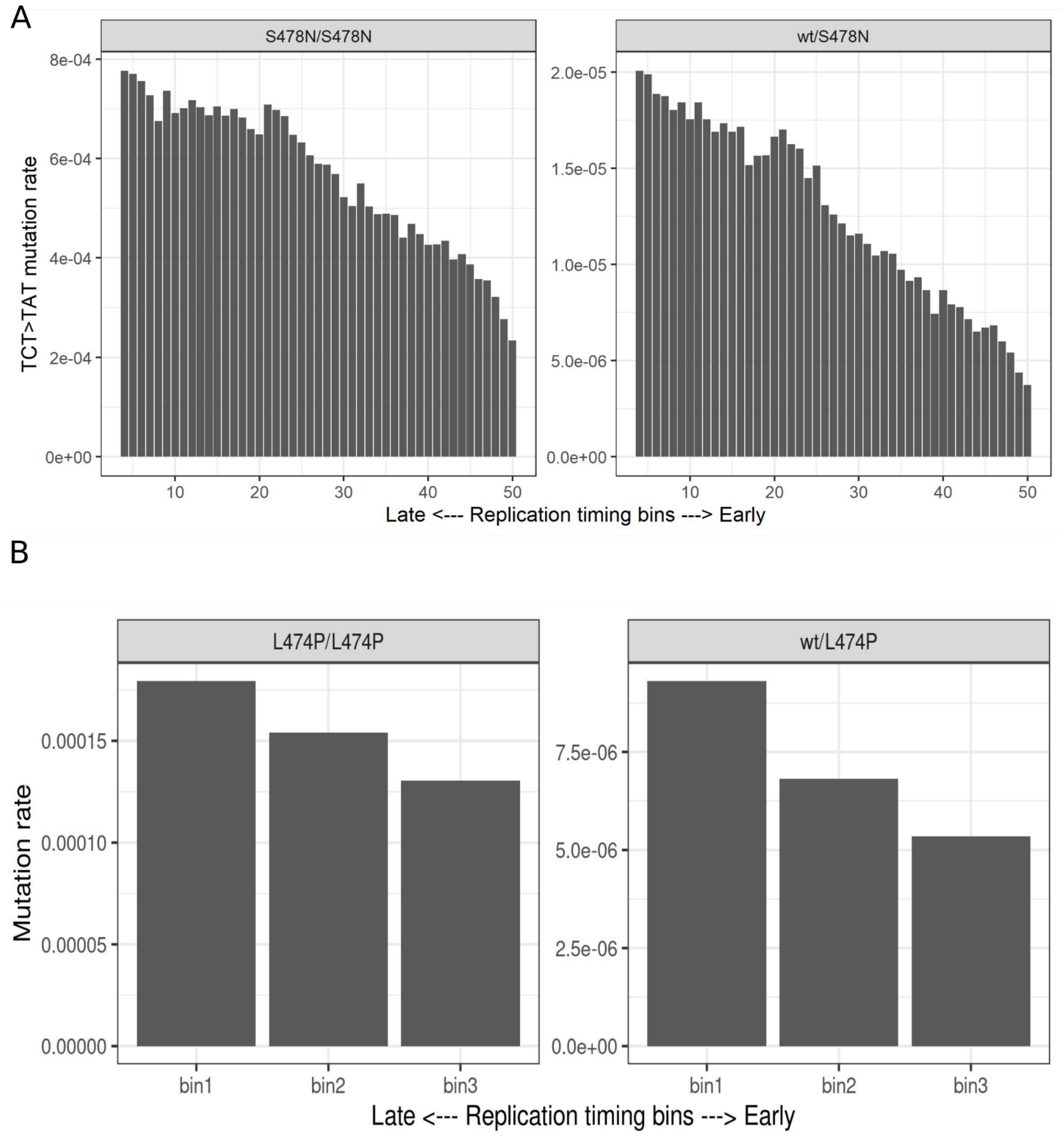


**Supplemental Figure S11.**

Dependence of mutation rate on replication timing in homozygous and heterozygous carriers of **A,** S478N (Robinson et al. 2021) and **B,** L474P (our data).

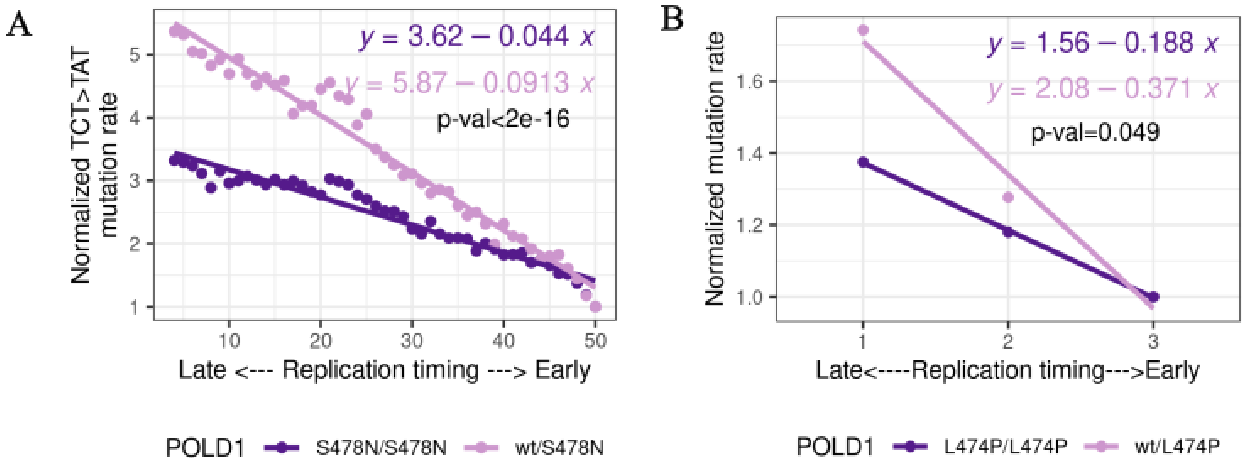


#### Supplemental Figure S12.

Dependence of mutation rate on replication timing in homozygous and heterozygous carriers of **A)** *POLD1* S478N ((Robinson et al. 2021)) and **B)** *POLD1* L474P (our data).

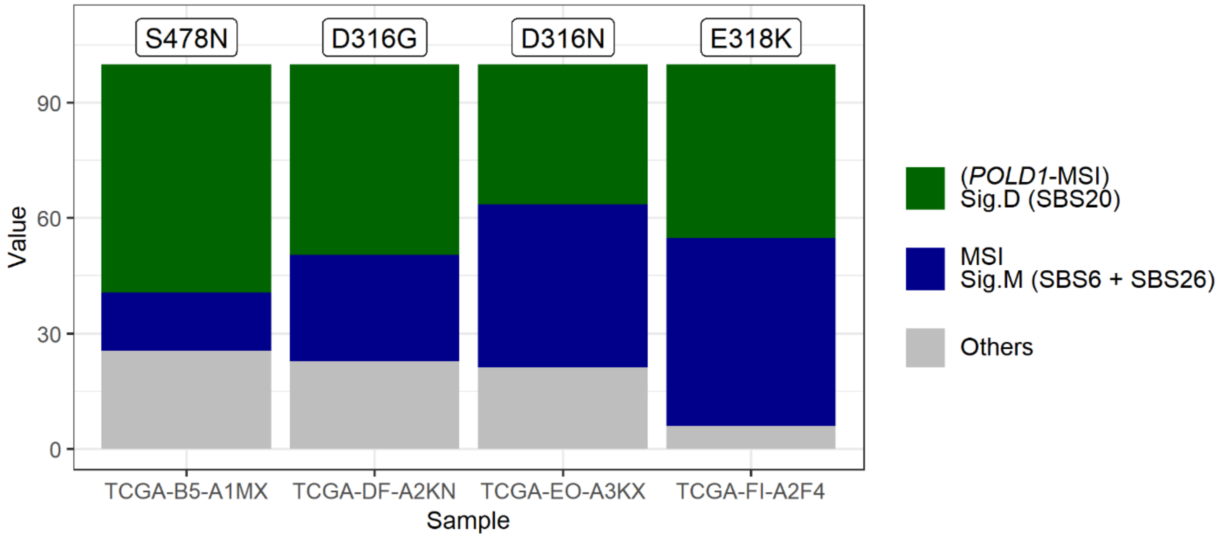


**Supplemental Figure S13.**

Mutational signatures in TCGA UCEC samples with somatic pathogenic variants in *POLD1*. Data obtained from previously published study (Haradhvala et al. 2018). Sig.D corresponding to simultaneous inactivation of MMR and *POLD1* proofreading is not an additive spectrum of MMR inactivation and POLD1 proofreading inactivation, thus it can be extracted separately from Sig.M with a high level of confidence.

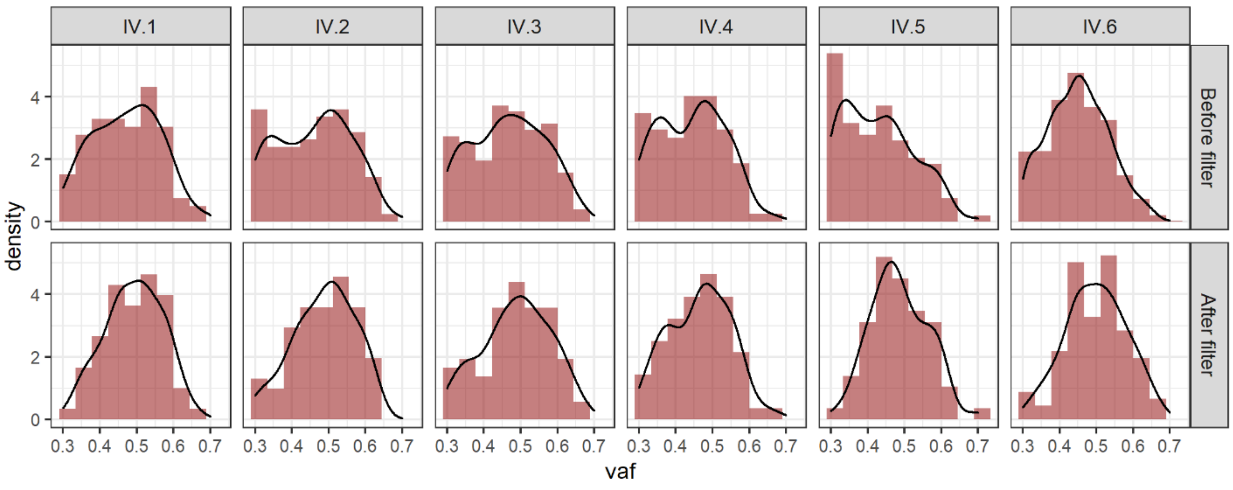


**Supplemental Figure S14.**

Variant allele frequency (VAF) distributions for candidate de novo mutations before and after filtering, using mutations identified in fibroblast colonies.

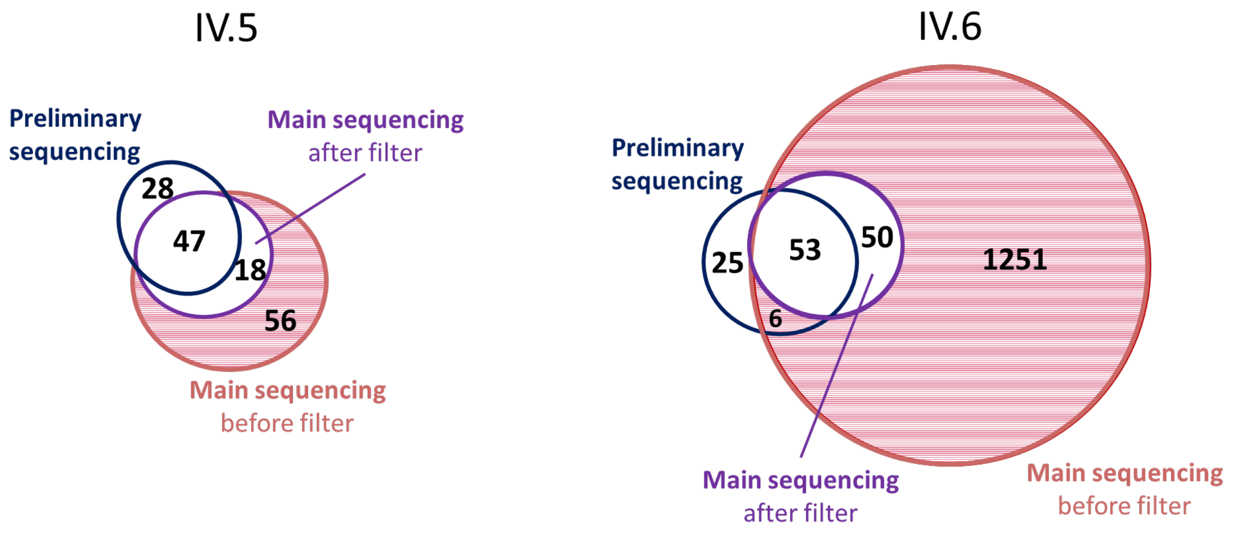


**Supplemental Figure S15.**

Intersection of candidate *de novo* mutations in preliminary sequencing of two trios and in main sequencing before and after the filtering for presence in the fibroblast colony was applied. Dashed red area corresponds to mutations removed by the filter.

**Supplemental Table S1.**

Clinical and phenotypic characteristics of the family members included in the study.

| Individual ID | Sex | *POLD1* status | Tissue sequenced for trio analysis | Single-cell colony from fibroblasts | MA experiment | Diagnosed Polyps/cancer (age of diagnosis) |
|---|---|---|---|---|---|---|
| IV.6 | Male | L474P | Fibroblasts | + | - | - |
| III.2 | Male | L474P | Fibroblasts | + | + | Endometrial cancer (58), 9 polyps (since age 48) |
| III.4 | Male | L474P | Blood | + | + | Colon cancer (50), >12 polyps (since age 50) |
| IV.4 | Female | L474P | Fibroblasts | + | + | 4 polyps (27) |
| III.6 | Male | L474P | Blood | - | - | Esophageal enteroid metaplasia, ~8 polyps (since age 48),<br>gastric fundic gland polyps (since age 57) |
| IV.2 | Female | L474P | Fibroblasts | + | + | 5-10 polyps (since age 25) |
| IV.1 | Female | L474P | Fibroblasts | + | - | Colon cancer (23), esophagus benign tumor* (30)<br>>50 gastric fundic gland polyps (since age 30) |
| IV.8 | Female | wt | Blood | - | - | - |
| III.7 | Female | wt | Blood | - | - | - |
| III.5 | Female | wt | Blood | - | - | - |
| IV.5 | Male | wt | Buccal swab | + | + | - |
| IV.3 | Male | wt | Fibroblasts | + | + | - |
| III.1 | Female | wt | Blood | - | - | - |
| III.3 | Female | wt | Blood | - | - | - |
| III.8 | Male | wt | Blood | - | - | - |
| IV.9 | Female | wt | Buccal swab | - | - | - |

*compatible with GIST by echoendoscopy.

Abbreviations: MA, mutation accumulation.

**Supplemental Table S2.**

Characteristics of the fibroblast cultures in the mutation accumulation experiment.

| Sample | Total # of passages | # of passages from single cell isolation until DNA sequencing | passage doubling rate | *POLD1* status |
|---|---|---|---|---|
| III.2 | 77 | 34 | 1.62 | L474P |
| IV.1 | 60 | 38 | NA | L474P |
| IV.2 | 87 | 46 | 1.58 | L474P |
| IV.3 | 67 | 35 | 1.49 | wt |
| III.4 | 60 | 38 | NA | L474P |
| IV.4 | 63 | 33 | 2.14 | L474P |
| IV.6 | 60 | 26 | NA | L474P |
| IV.5 | 54 | 33 | 2.07 | wt |

#### Supplemental Table S3.

Statistical significance of presence of SBS10c signature in mutations accumulated during the experiment calculated using mSigAct package.

| Sample | *POLD1* status | p-value |
|---|---|---|
| IV.6 | L474P | 1.430436e-24*** |
| III.2 | L474P | 1.817e-34*** |
| IV.5 | wt | 0.01964926* |
| III.4 | L474P | 8.094666e-24** |
| IV.3 | wt | 0.8072493 |
| IV.4 | L474P | 3.63662e-55** |

#### Supplemental Table S4.

Probability of LOH in the studied samples.

| | Sites with cnLOH | Sites with LOH (cnLOH or deletion) | Target sites | cnLOH probability | LOH probability |
|---|---|---|---|---|---|
| L474P | 3036874 | 3603390 | 2628131683 | $1.16*10^{-3}$ | $1.37*10^{-3}$ |
| D316H | 143778197 | 266663050 | 2587827238 | $5.56*10^{-2}$ | $1.03*10^{-1}$ |
| S478N | 661469 | 18389853 | 2615521263 | $2.53*10^{-4}$ | $7.03*10^{-3}$ |
| Simultaneous probability | | | | $1.63*10^{-8}$ | $9.92*10^{-7}$ |

**Supplemental Table S5.**

MMR status of tumors with inherited *POLD1* pathogenic mutations.

| *POLD1* nucleotide substitution | *POLD1* amino acid substitution | Cancer type | Reference | MSI/MSS status |
|---|---|---|---|---|
| c.947A>G | D316G | Colorectal | Bellido et al. 2016 | MSS |
| c.947A>G | D316G | Endometrial | Bellido et al. 2016 | MSS |
| c.1433G>A | S478N | Colorectal | Palles et al. 2013 and 2021 | MSS<br>(tumor, 1 AP) |
| c.1433G>A | S478N | Adenoma | Palles et al. 2013 and 2021 | MSS<br>(5 AP) |
| c.1433G>A | S478N | Endometrial | Palles et al. 2013 and 2021 | MSS |
| c.1433G>A | S478N | Colorectal | Ito et al. 2020 | Normal MMR protein expression |
| c.1421T>C | L474P | Colorectal | Valle et al. 2014 | MSS |
| c.1421T>C | L474P | Colorectal | Bellido et al. 2016 | MSS |
| c.1421T>C | L474P | Colorectal | Ferrer-Avargues et al. 2017 | MSI |
| c.1421T>C | L474P | Colorectal | Ferrer-Avargues et al. 2017 | MSS |

Abbreviations: AP, adenomatous polyp; MMR, DNA mismatch repair; MSI, microsatellite instability; MSS, microsatellite stability
